# A live attenuated vaccine confers superior mucosal and systemic immunity to SARS-CoV-2 variants

**DOI:** 10.1101/2022.05.16.492138

**Authors:** Geraldine Nouailles, Julia M. Adler, Peter Pennitz, Stefan Peidli, Gustavo Teixeira Alves, Morris Baumgart, Judith Bushe, Anne Voss, Alina Langenhagen, Fabian Pott, Julia Kazmierski, Cengiz Goekeri, Szandor Simmons, Na Xing, Christine Langner, Ricardo Martin Vidal, Azza Abdelgawad, Susanne Herwig, Günter Cichon, Daniela Niemeyer, Christian Drosten, Christine Goffinet, Markus Landthaler, Nils Blüthgen, Haibo Wu, Martin Witzenrath, Achim D. Gruber, Samantha D. Praktiknjo, Nikolaus Osterrieder, Emanuel Wyler, Dusan Kunec, Jakob Trimpert

## Abstract

Vaccines are a cornerstone in COVID-19 pandemic management. Here, we compare immune responses to and preclinical efficacy of the mRNA vaccine BNT162b2, an adenovirus-vectored spike vaccine, and the live-attenuated-virus vaccine candidate sCPD9 after single and double vaccination in Syrian hamsters. All regimens containing sCPD9 showed superior efficacy. The robust immunity elicited by sCPD9 was evident in a wide range of immune parameters after challenge with heterologous SARS-CoV-2 including rapid viral clearance, reduced tissue damage, fast differentiation of pre-plasmablasts, strong systemic and mucosal humoral responses, and rapid recall of memory T cells from lung tissue. Our results demonstrate that use of live-attenuated vaccines may offer advantages over available COVID-19 vaccines, specifically when applied as booster, and may provide a solution for containment of the COVID-19 pandemic.

## Introduction

As of early 2022, ten COVID-19 vaccines fulfill the quality, safety and efficacy requirements that allow emergency use listing (EUL) by the WHO (*1*). Currently authorized vaccines employ classical approaches such as inactivated virus and subunit vaccines, as well as more recent technology, including the use of adenoviral vectors and the novel nucleoside-modified mRNA vaccines (*2*). Despite initially high vaccine efficacy and long-lasting protection from severe illness, waning of protection from infection and symptomatic disease is now evident, particularly following the emergence and spread of the omicron variant (*3*, *4*). Moreover, global disparity in vaccine access remains alarmingly high: By 6 May 2022, only 15.8% of people in low-income countries had received at least one dose of vaccine despite the surplus possessed by wealthy nations (*5*). This situation, together with the continued evolution of SARS-CoV-2, warrants an even more robust vaccine and immunization strategy. Optimal COVID-19 vaccines would not just protect from severe disease, but also provide protection from infection with a broad spectrum of virus variants, while, at the same time, they would prevent or significantly limit SARS-CoV-2 transmission. Although live attenuated vaccines (LAV) are used very successfully to tackle many virus infections such as measles, mumps and rubella (MMR) (*6*), studies investigating the efficacy, effectiveness and cross-comparison of promising intranasal LAVs with other routes of delivery remain limited (*7*, *8*). Notably, LAVs do not depend on adjuvants (*7*) and can be administered locally, for example intranasally, as is the case for influenza LAVs (*9*). Comprised of replication-competent viruses, intranasal LAVs mimic the natural course of infection and antigen production, which distinguishes them from locally administered, replication-incompetent vector- or antigen-based vaccines (*10*). In contrast to empirically generated vaccines used in the past, modern LAV design utilizes molecular tools to limit virus replication and virulence, while maintaining immunogenicity and antigenic integrity (*11*). One recent strategy employed in rational design of LAVs is codon pair deoptimization (CPD), suitable for both DNA (*12*, *13*) and RNA viruses (*14*, *15*), including SARS-CoV-2 (*16*).

Currently approved COVID-19 vaccines are administered intramuscularly and can efficiently induce protective systemic immunity, including high titers of neutralizing serum antibodies, central and effector memory T cells (*17*), germinal center B cells (*18*) and long-lived plasma cells (LLPC) (*19*). Yet, the vaccines are less efficient in inducing durable mucosal IgA and IgG responses (*20*–*22*) as well as pulmonary tissue-resident memory cell responses (*23*). Furthermore, pathogen-specific mucosal antibodies at the site of virus entry are considered central to limiting infectivity and transmission (*24*). Accordingly, tissue-resident memory cells undergo faster recall responses, as their local positioning allows for earlier cognate antigen recognition (*25*). Hence, vaccines effectively administered via the respiratory entry route, are expected to induce robust local mucosal immunity against the targeted pathogen (*26*, *27*).

In this study, we sought to compare different approaches to vaccination and evaluate potential differences in systemic and mucosal immunity conferred by different vaccines as well as a various prime-boost vaccine regimens that included systemic priming followed by a respiratory boost.

## Methods

### Study design

This study compared the protective potential of the modified live attenuated SARS-CoV-2 mutant sCPD9 (*28*, *30*), adenovirus vector vaccine candidate Ad2-spike (*28*) and mRNA vaccine BNT162b2. Protection was assessed in Syrian hamsters against challenge infection with the SARS-CoV-2 Delta variant. The study also employs single cell sequencing to characterize the innate and adaptive immune response to vaccination and challenge infection with virulent SARS-CoV-2. Syrian hamsters were vaccinated with either one (prime only experiment) or two doses (prime-boost experiment) of the three vaccines studied. Inoculation was performed either intranasally with the modified live attenuated virus sCPD9 or intramuscularly with the Ad2-spike or mRNA vaccine. In the prime only experiment, hamsters were challenge-infected with SARS-CoV-2 Delta variant (1 × 10^5^ plaque-forming units (PFU), 60 μL) by intranasal instillation 21 days post vaccination. In the prime-boost experiment, hamsters were boosted with a second vaccine dose on day 21 following the prime dose. 14 days post booster vaccination (35 days post prime), hamsters were challenge-infected with SARS-CoV-2 Delta variant (1 × 10^5^ PFU, 60 μL) by intranasal instillation. At day 2 and 5 after challenge, hamsters were euthanized, and blood and parts of the upper and lower airways (pharynx, trachea, and lungs) were collected for viral titrations, RT-qPCR, histopathological examinations, and single-cell sequencing.

### Cells

Vero E6 (ATCC CRL-1586) and VeroE6-TMPRSS2 (NIBSC 100978) cells were cultured in minimal essential medium (MEM) containing 10 % fetal bovine serum, 100 IU/mL penicillin G, and 100 μg/mL streptomycin at 37°C and 5 % CO_2_. In addition, the cell culture medium for Vero-TMPRSS2 cells contained 1000 μg/mL geneticin (G418) to ensure selection for cells expressing the genes for neomycin resistance and TMPRSS2.

### Viruses

The modified live attenuated SARS-CoV-2 mutant sCPD9 and SARS-CoV-2 variants B.1 - BetaCoV/Munich/ChVir984/2020 (B.1, EPI_ISL_406862), Beta – B.1.351 (hCoV-19/Netherlands/NoordHolland_20159/2021), and Delta – B.1.617.2 (SARS-CoV-2, Human, 2021, Germany ex India, 20A/452R (B.1.617) were propagated on VeroE6-TMPRSS2 cells. Omicron BA.1 - B.1.1.529.1 (hCoV-19/Germany/BE-ChVir26335/2021, EPI_ISL_7019047) was propagated on CaLu-3 cells. Prior to experimental infection virus stocks were stored at −80°C.

### Ethics statement

*In vitro* and animal work was conducted under the appropriate biosafety conditions in the biosafety level three (BSL-3) facility at the Institut für Virologie, Freie Universität Berlin, Germany. All animal experiments were performed in compliance with relevant institutional, national, and international guidelines for care and humane use of animal and approved by the Landesamt für Gesundheit und Soziales in Berlin, Germany (permit number 0086/20).

### Animal Husbandry

Nine- to eleven-week-old Syrian hamsters (*Mesocricetus auratus;* breed RjHan:AURA) were purchased form Janvier Labs and were housed in groups of 2 to 3 animals in individually ventilated cages (IVCs). The hamsters had free access to food and water. They were allowed to get used to the housing conditions for seven days prior to vaccination. For both experiments, the cage temperatures were constantly between 22 and 24°C with a relative humidity between 40 and 55%.

### Infection Experiments

We studied the efficacy of the BNT162b2 mRNA vaccine (Pfizer-BioNTech), adenovirus vector vaccine candidate Ad2-spike (*28*) and modified live attenuated SARS-CoV-2 vaccine candidate sCPD9 (*29*, *30*), as well as immune responses to vaccination and challenge in two consecutive and independent animal experiments. The infection experiments were done in the Syrian hamsters, a highly susceptible model for SARS-CoV-2 infection (*70*). Hamsters were randomly assigned into groups, with 50 – 60 % of the animals in each group being female.

In the first experiment, 15 hamsters were mock-vaccinated or vaccinated with live attenuated sCPD9 virus, Ad2-spike or mRNA. Vaccination with sCPD9 was done by intranasal instillation under anesthesia (1 × 10^5^ focus-forming units (FFU), 60 μL) (*70*), Ad2-spike (5 × 10^8^ infectious units, 200 μL), and mRNA vaccine (5 μg mRNA, 50 μL) by intramuscular injection. Mock-vaccinated hamsters were vaccinated by intranasal instillation with sterile cell culture supernatant obtained from uninfected VeroE6-TMPRSS cells. 21 days after vaccination, hamsters were challenge-infected with SARS-CoV-2 Delta variant (1 × 10^5^ PFU, 60 μL) by intranasal instillation under anesthesia. In the second experiment, 10 hamsters were either mock-vaccinated or vaccinated with one of the three vaccines (see above) followed by a booster vaccination 21 days later with the same or a different vaccine. 14 days after booster vaccination, the hamsters were challenged as described above.

Infected hamsters were inspected twice daily for clinical signs of infection. In the first experiment, body weights were recorded daily for the duration of the experiment. In the second experiment, body weights were recorded only after challenge infection, because we had confirmed in the first animal experiment that all vaccines were safe for hamsters and had no effect on body weight development.

In the first animal experiment, 5 hamsters from each group were euthanized on day 21 after vaccination and days 2 and 5 after challenge (days 23 and 26 of the experiment). In the second animal experiment, blood was taken from all animals via saphenous vein puncture under anesthesia on day 35 prior to infection. 5 hamsters from each group were euthanized on days 2 and 5 after challenge (day 37 and 40 of the experiment).

Blood, tracheal swabs, and portions of the airway were collected to determine virological, histological, and molecular parameters of vaccination and immune response. The left lung was preserved in 4 % formaldehyde solution for detailed histopathological examination.

### Vaccine preparations

sCPD9 was grown on Vero-TMRSS cells and titrated on Vero E6 cells as described previously, final titers were adjusted to 2 × 10^6^ FFU/mL in MEM. Recombinant Ad2-spike was generated, produced on 293 T cells and purified as previously described. BNT162b2 was obtained as commercial product (Comirnaty^®^) and handled exactly as recommended by the manufacturer with the exception that the final concentration of mRNA was adjusted to 50 μg/mL (100 μg/mL is the recommended concentration for use in humans) by adding injection grade saline (0.9 % NaCl in sterile water) immediately prior to use.

### Vaccination

sCPD9 was applied intranasally under general anesthesia (0.15 mg/kg medetomidine, 2.0 mg/kg midazolam and butorphanol 2.5 mg/kg) at a dose of 1 × 10^5^ FFU per animal in a total volume of 60 μL MEM. Ad2-spike was injected intramuscularly at 5 × 10^8^ infectious units in 200 μL injection buffer (3 mM KCl, 1 mM MgCl2, 10 % glycerol in PBS). BNT162b2 was injected intramuscularly at 5 μg per animal in 100 μL physiological saline (0.9 % NaCl in sterile water).

### Nasal washes

Nasal washes were obtained from each hamster in this study. To this end, the skull of each animal was split slightly paramedian, such that the nasal septum remained intact on one side of the nose. Subsequently, a 200 μL pipette tip was carefully slid underneath the nasal septum and 150 μL wash fluid (PBS with 30 μg/mL ofloxacin and 10 μg/mL voriconazole) was applied. The wash was collected through the nostril and the washing procedure was repeated twice, approximately 100 μL of sample was recovered after the third wash.

Nasal washes obtained from the prime-only vaccination trial were subjected to ELISA analysis of SARS-CoV-2 Spike specific IgA antibodies. Nasal washes obtained from the prime-boost vaccination trial were used for microneutralization assay to assess their capacity to neutralize the SARS-CoV-2 ancestral variant B1.

### Plaque assay

For quantification of replication-competent virus, 50 mg of lung tissue were used. Serial 10-fold dilutions were prepared after homogenizing the organ samples in a bead mill (Analytic Jena). The dilutions were plated on Vero E6 cells grown in 24-well plates and incubated for 2.5 h at 37°C. Subsequently, cells were overlaid with MEM containing 1.5 % carboxymethylcellulose sodium (Sigma Aldrich) and fixed with 4 % formaldehyde solution 72 hours after infection. To count the plaque-forming units, plates were stained with 0.75 % methylene blue.

### Histopathology, immunohistochemistry and in situ-hybridization

Lungs were processed as previously described (*70*). After careful removal of the left lung lobe, tissue was fixed in PBS-buffered 4 % formaldehyde solution, pH 7.0 for 48 h. For conchae preparation, parts of the left skull half were fixed accordingly. Afterwards, lungs or conchae were gently removed from the nasal cavity and embedded in paraffin, cut at 2 μm thickness and stained with hematoxylin and eosin (H&E).

In situ-hybridization on lungs was performed as previously described (*71*) using the ViewRNA™ ISH Tissue Assay Kit (Invitrogen by Thermo Fisher Scientific, Darmstadt, Germany) according to the manufacturer’s instructions with minor adjustments. For SARS-CoV-2 RNA localization, probes detecting N gene sequences (NCBI database NC_045512.2, nucleotides 28,274–29,533, assay ID: VPNKRHM) were used. Sequence-specific binding was controlled by using a probe for detection of pneumolysin. Immunohistochemistry (IHC) on conchae was performed as described earlier (*72*). Paraffin-embedded tissues were cut (2 μm thickness), mounted on adhesive glass slides, dewaxed in xylene, rehydrated in descending grades of alcohol, and endogenous peroxidase was inhibited. Antigen retrieval was performed using microwave heating (600 W) in 10 mM citric acid (pH 6.0) for 12 min for SARS-CoV-1 nucleoprotein antibody (Sino Biological Inc.; Beijing, China) and using recombinant protease from *Streptomyces griseus* (PanReac Applichem, Darmstadt, Germany) for 13 min at 37°C for IgA antibody. For blockage of non-specific antibody binding incubation with 8 % Roti-Immunoblock (Roth, Karlsruhe, Germany) and 20 % normal goat serum for 30 min was implemented. Anti-SARS-CoV-1 NP mouse monoclonal antibody (Sino Biological Inc.; Beijing, China, dilution: 1:500) and rabbit anti hamster IgA antibody (Brookwood Biomedical, Jemison, AL, dilution: 1:250) were incubated at 4°C overnight followed by washing and incubation with a secondary biotinylated goat anti-mouse IgG antibody (dilution: 1:200, Vector Laboratories, Burlingame, California, USA). For color development freshly prepared avidin-biotin-peroxidase complex (ABC) solution (Vectastain Elite ABC Kit; Vector Laboratories) was incubated for 30 min followed by incubation with diaminobenzidine tetrahydrochloride (Merck, Darmstadt, Germany) for 4 min. Slides were counterstained with Mayer’s hematoxylin.

Blinded microscopic analysis was performed by board-certified veterinary pathologists (JB). An Olympus BX41 microscope with a DP80 Microscope Digital Camera and the cellSensTM Imaging Software, Version 1.18 (Olympus Corporation, Münster, Germany) was utilized for histopathological evaluations and photographs. Slides were automatically digitized with the Aperio CS2 slide scanner (Leica Biosystems Imaging Inc., Vista, CA, USA) and overviews were generated by using the image Scope Software (Leica Biosystems Imaging Inc.).

### Neutralization assays from nasal washes

To assess the capacity of nasal washes obtained from the prime-boost vaccination trial with respect to neutralization of authentic SARS-CoV-2 (B.1), nasal washes were diluted 1:1 in 2× MEM containing 50 μg/mL enrofloxacin and 10 μg/mL voriconazole, subsequent serial dilutions were performed in MEM containing 25 mg/mL enrofloxacin, 5 μg/mL voriconazole and 1 % FBS. SARS-CoV-2 (50 PFU) were added to the nasal wash dilutions and dilutions from 1:2 to 1:256 were plated on near-confluent Vero E6 cells seeded in 96-well cell culture plates. At 3 days after inoculation, cells were fixed and stained with methylene blue. To identify virus-neutralizing dilutions, the integrity of the cell monolayer was assessed by comparison with control wells that contained either no nasal wash or no virus. The last dilution in which no evidence for virus-induced cpe was seeable was considered the neutralizing titer for the respective sample.

### Enzyme-linked Immunosorbent Assays (ELISA) from nasal washes

An in-house ELISA was performed to investigate SARS-specific IgA levels in nasal washes after vaccination. MEDISORP plates (Thermofisher, MW96F straight) were used to conduct this assay and coated in two steps. First, SARS-CoV-2 spike protein (Acro Biosystems, SARS-CoV-2 S protein (D614G), His Tag, Super stable trimer) was diluted in 1× PBS to a final concentration of 20 μg/mL, and 5 μL of the respective dilution were applied to each well. The antigen was diluted in tubes with a low binding capacity for proteins (Eppendorf Protein LoBind Tube 1.5 ml). Second, 45 μL of coating buffer (diH2O + 5.3 g Na2CO3 (50 mM) + anhydrous 4.2 g NaHCO3 (50 mM), pH 9.6) were added per well, followed by an incubation time of 12 h at 4°C. Subsequently, the plates were washed four times with washing buffer (1 × PBS + 0.05 % Tween20) and blocked with 1× PBS + 1 % BSA + 10 % FCS for 1 h. Nasal washes (each sample in duplicate) were diluted 1:100 in dilution buffer (1 × PBS + 2 % BSA + 0.1 % Tween 20) and 50 μL were pipetted to each well. The plates were covered and incubated for 2 h at room temperature before the washing step was repeated and 50 μL of secondary antibody (Brookwood biomedical, Rabbit Anti-Hamster IgA, HRP-conjugated) diluted 1:1000 was applied per well. The covered plates were then incubated for 1 h at room temperature and washed again as described above. Subsequently, 50 μL of 3,3’,5,5’-Tetramethylbenzidine (TMB) were added followed by another 15 min of incubation. After applying 50 μL stop solution (1 M H2SO4) per well, SpectraMax Plus384 was used to read the plates at 450 nm and 570 nm to assess optical density (OD).

### Serum neutralization assay

This test was conducted to determine the neutralizing activity against SARS-CoV-2 (B.1) and different variants of concern (Beta, Delta, Omicron) of serum samples collected in the prime and the prime-boost experiment. Day 0 samples of the prime-boost trial could not be tested for neutralizing antibodies against B.1.351 (Beta) due to lack of material. Sera were inactivated at 56°C for 30 min. Two-fold serial dilutions (1:8 to 1:1024) were plated on 96-well plates and 200 PFU SARS-CoV-2 were pipetted into each well. After an incubation time of 1 h at 37°C the dilutions were transferred to 96-well plates containing sub-confluent Vero E6 cells and incubated for 72 h at 37°C (B.1, Beta, Delta) or for 96 h at 37°C (Omicron). The plates were fixed with 4 % formaldehyde solution and stained with 0.75 % methylene blue. Wells that showed no cytopathic effect were considered neutralized.

### RNA extraction and qPCR

To quantify genomic copies in oropharyngeal swabs and 25 mg homogenized lung tissue, RNA was extracted using innuPREP Virus DNA/RNA Kit (Analytic Jena, Jena, Germany) according to the manufacturer’s instructions. The qPCR was performed using the NEB Luna Universal Probe One-Step RT-qPCR Kit (New England Biolabs) with cycling conditions of 10 min at 55°C for reverse transcription, 3 min at 94°C for activation of the enzyme, and 40 cycles of 15 s at 94°C and 30 s at 58°C on a qTower G3 cycler (Analytic Jena, Jena, Germany) in sealed qPCR 96-well plate. Primers and probes were used as previously reported (*73*).

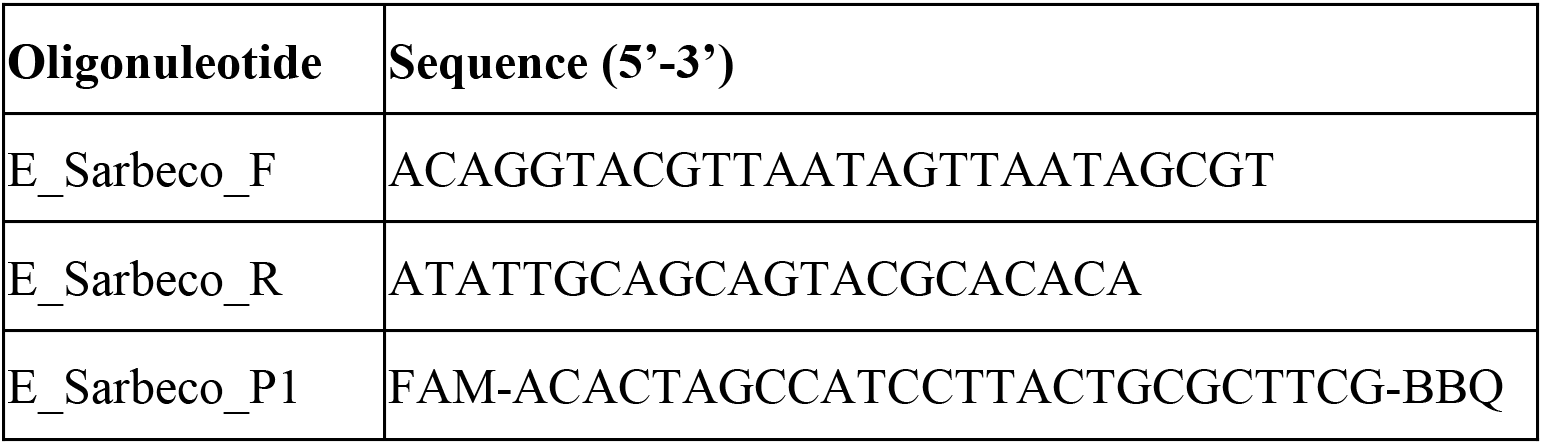

### *Mesocricetus auratus* genome annotation

For quantification of gene expression, we used the MesAur 2.0 genome assembly and annotation available via the NCBI genome database (https://www.ncbi.nlm.nih.gov/genome/11998?genome_assembly_id=1585474). The GFF file was converted to GTF using gffread (74). Where no overlaps were produced, 3’-UTRs in the annotation were extended by 1000 bp as described previously (75). Further polishing steps for the GTF file are described on the github page accompanying this manuscript (https://github.com/Berlin-Hamster-Single-Cell-Consortium/Live-attenuated-vaccine-strategy-confers-superior-mucosal-and-systemic-immunity-to-SARS-CoV-2). The final gtf file used for the analysis is available through GEO (https://www.ncbi.nlm.nih.gov/geo/query/acc.cgi?acc=GSE200596).

### Bulk RNA extraction

To perform RNA bulk sequencing, RNA was isolated from lung tissue using Trizol reagent according to the manufacturer’s instructions (ambion, life technologies). Briefly, 1 ml Trizol was added to the homogenized organ sample and vortexed thoroughly. After an incubation time of 20 min, 200 μL of chloroform were added. The samples were vortexed again and incubated for 10 min at room temperature. Subsequently, tubes were centrifuged at 12,000 × *g* for 15 min at 4°C and 500 μL of the aqueous phase were transferred into a new tube containing 10 μg GlycoBlue. Isopropanol (500 μL) was added followed by vortexing, incubating and centrifuging the samples as described above. Thereafter, isopropanol was removed and 1 mL of ethanol (75 %) was applied. The tubes were inverted shortly and centrifuged at 8.000× *g* for 10 min. After freeing the pellet from ethanol, RNA was resuspended in 30 μL of RNAase-free water and stored at −80°C.

### Cell isolation from blood and lungs

White blood cells were isolated from EDTA-blood as previously described, steps included erylysis and cell filtration prior counting. Lung cells (caudal lobe) were isolated as previously described (*32*), steps included enzymatic digestion, mechanical dissociation and filtration prior counting in trypan blue. Buffers contained 2 μg/mL actinomycin D to prevent *de novo* transcription during the procedures.

### Cell isolation from nasal cavities

To obtain single cell suspensions from the nasal mucosa of SARS-CoV-2 challenged hamsters, the skull of each animal was split slightly paramedian, such a way that the nasal septum remained intact on the left side of the nose. The right side of the nose was carefully removed from the cranium and stored in ice-cold 1× PBS with 1 % BSA and 2 μg/mL actinomycin D until further use. Nose parts were transferred into 5 mL Corning^®^ Dispase solution supplemented with 750 U/mL Collagenase CLS II, and 1 mg/mL DNase and incubated at 37°C for 15 min. For preparation of cells from the nasal mucosa, the conchae were carefully removed from the nasal cavity and re-incubated in digestion medium for 20 min at 37°C. Conchae tissue was dissociated by pipetting and pressing through a 70-μm filter with a plunger. Ice-cold PBS with 1 % BSA and 2 μg/mL actinomycinD was added to stop the enzymatic digestion. Cell suspension was centrifuged at 4°C for 15 min at 400× *g* and supernatant discarded. The pelleted nasal cells were resuspended in 5 mL red blood cell lysis buffer and incubated at RT for 2 min. Lysis reaction was stopped with 1 × PBS with 0.04 % BSA and cells centrifuged at 4°C for 10 min at 400× *g*. Pelleted cells were resuspended in 1× PBS with 0.04 % BSA and 40 μm-filtered. Live cells were counted in trypan blue and viability rates determined using a counting chamber. Cell concentration for scRNA-seq was adjusted by dilution.

### Single-cell RNA sequencing

Isolated cells from blood, lungs and nasal cavities of Syrian hamsters were subjected to scRNA-Seq using the lO× Genomics Chromium Single Cell 3’ Gene Expression system with Feature Barcoding technology for Cell Multiplexing. In the prime only experiment, ~ 500,000 cells per blood sample, 1,000,000 cells per lung sample, and 1,000,000 per nasal cavity sample underwent cell multiplex oligo (CMO) labeling according to manufacturers’ instructions (3’ CellPlex Kit Set A; 10× Genomics), using the classical labelling protocol. In the prime-boost experiment, ~150,000 cells per blood sample, ~350,000 cells per lung sample, and ~350,000 per nasal cavity sample underwent cell multiplex oligo (CMO) labeling according to manufacturers’ instructions (3’ CellPlex Kit Set A; 10× Genomics), using the 96-well plate labelling protocol for low cell numbers to minimize protocol duration and cell manipulation (https://assets.ctfassets.net/an68im79xiti/4G3MABhAGLG8oAEyrSiz7L/a6e4f846634a50f0d44_025bab4b0f858/CG000426_PlateBasedSamplePrep_RevC.pdf.). In the first two experimental runs (prime experiment), pools were generated from 12 samples derived from all three organ types. De-multiplexing difficulties for blood samples from mixed pools led to a change in protocol for the third and fourth experimental run (prime-boost experiment). Here, multiplex pools were generated from 8 samples of the same organ origin only.

Oligo labelled cells were pooled accordingly without prior counting to minimize protocol duration, assuming equal cell loss between samples during the labelling procedures. Pooled cells were 40-μm filtered, counted and cell concentration adjusted to 1450 −1,600 cells /μL, resulting in loading of 49,500 pooled cells per lane aiming to recover 30,000 cells. Pools consisting of 12 samples were loaded onto 4 Chromium Next GEM Chip G lanes, pools consisting of 8 samples were loaded onto 2 or 3 lanes (blood samples on 2 lanes, lung and nasal cavity on 3 lanes). Loaded cells underwent partitioning, barcoding and mRNA reverse transcription in Gel-Beads-in-Emulsions following the instructions of Chromium Next GEM Single Cell 3’ Reagent Kits v.3.1 (Dual Index) provided by the manufacturer (10× Genomics). Viability rates of nasal cavity cells did not allow for qualitative results for the prime-boost experiment. Library sequencing was performed on a Novaseq 6000 device (Illumina), with SP4 flow cells (read1: 28 nt; read2: 150 nt).

### Analysis of single-cell RNA sequencing data

Sequencing reads were initially processed using the multi command of the 10× Genomics Cell Ranger 6.0.2 software. For the cellplex demultiplexing, the assignment thresholds were partially adjusted (see the github page accompanying this manuscript, https://github.com/Berlin-Hamster-Single-Cell-Consortium/Live-attenuated-vaccine-strategy-confers-superior-mucosal-and-systemic-immunity-to-SARS-CoV-2, for details). Further processing was done using the Seurat R package (76). In the next step, cells were filtered by a loose quality threshold (minimum 250 detected genes per cell) and clustered. Cell types were then annotated per cluster, and filtered using cell type specific thresholds (cells below the median or in the lowest quartile within a cell type were removed). The remaining cells were processed using the SCT/integrate workflow (77), and cell types again annotated on the resulting Seurat object. All code for downstream analysis is available on github, https://github.com/Berlin-Hamster-Single-Cell-Consortium/Live-attenuated-vaccine-strategy-confers-superior-mucosal-and-systemic-immunity-to-SARS-CoV-2.

## Results

Utilizing the widely employed mRNA vaccine BNT162b2 and two original vaccine candidates – an adenoviral vector vaccine carrying the spike glycoprotein of SARS-CoV-2 (*28*) and a live attenuated COVID-19 vaccine candidate named sCPD9 (*29*, *30*) – we assessed vaccine efficacy and mode of action in a heterologous SARS-CoV-2 Delta variant challenge setting.

First, Syrian hamsters were vaccinated with a single dose (prime-only regimen) of any of the three vaccines mentioned above and challenged with the SARS-CoV-2 Delta variant 21 days post-vaccination. In a second experimental setup, hamsters were immunized with two vaccine doses (prime-boost regimen) in which the booster was applied 21 days after the initial vaccination and followed by Delta variant challenge 14 days after the boost (Figure 1A). All vaccines were well tolerated, no signs of adverse effects were observed in the 21 days following the initial vaccination, and hamsters steadily gained body weight during this time (Figure S1A).

**FIG. 1.**
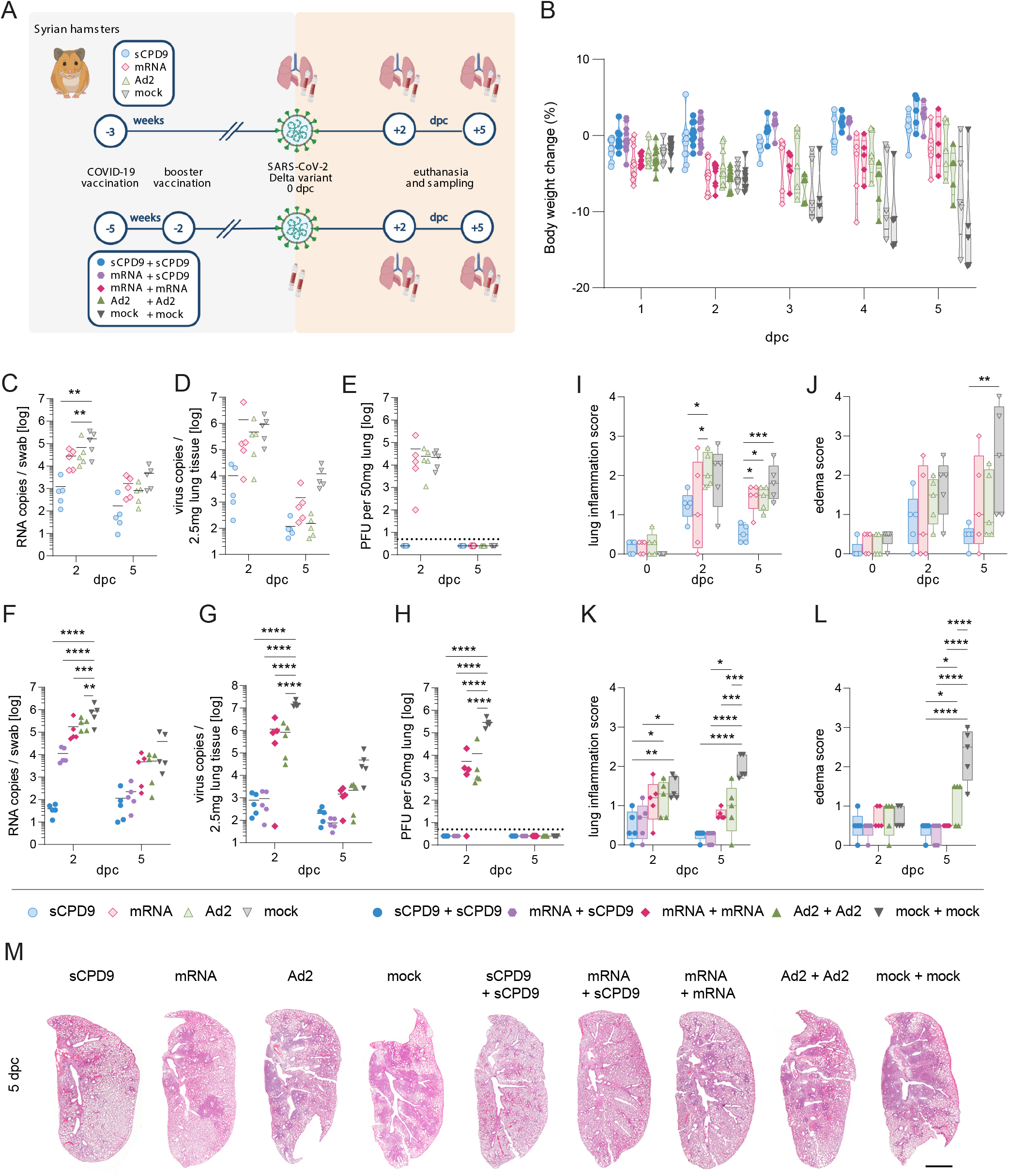
Clinical, virological and pathological parameters of SARS-CoV-2 infection in vaccinated and non-vacci-nated hamsters. **(A)** Experimental scheme showing setup and timelines for prime and prime-boost vaccinations. Syrian hamsters were vaccinated as indicated and challenged with SARS-CoV-2 (1 × 10^5^ pfu SARS-CoV-2 Delta). **(B)** Body weight development in percent after virus challenge were measured until analysis time point and displayed according to vaccination group. Violine plot (truncated) with quartiles and median. **(C – E, I – J)** Results of prime vaccinated and challenged animals and **(F – H, K – L)** results of prime-boost vaccinated and challenged animals: Number of genomic RNA (gRNA) copies detected in oropharyngeal swabs **(C, F)** and homogenized lung tissue **(D, G)**. **(E, H)** Quantification of replication-competent virus as plaque-forming units (pfu) per 50 mg homogenized lung tissue is shown. Dotted line marks the limit of detection (DL = 5 pfu). Titers below the detection limits were set to DL/2 = 2.5 pfu. **(I, K)** Lung inflammation was scored including severity of pneumonia, alveolar epithelial necrosis and endothelialitis. **(J, L)** Lung edema score accounting for perivascular and alveolar edema is displayed. **(M)** Haematoxylin-eosin-stained left lung sections illustrate different severities of pneumonia including peribronchial cuffs and consolidated areas between different vaccine schedules and non-vaccinated animals at 5 dpc. **(C – H)**, Scatter dot blots, line indicates mean, symbols represent individual hamsters. **(I – L)**, Box, 25th to 75th percentile and whiskers, Min to Max, symbol represent individual hamsters. **(C – L)**, Two-way ANOVA and Tukey’s multiple comparisons test are shown. * p < 0.05, ** p < 0.01, *** p < 0.001, and **** p < 0.0001.

### Vaccination alleviates clinical symptoms and reduces virus load

All vaccines used here protected animals from the considerable body weight loss that is typical for SARS-CoV-2 infection of Syrian hamsters (Figure 1B). At the same time, vaccine regimens involving sCPD9 displayed a trend towards improved protection compared to other vaccines based on body weight loss (Figure 1B).

Following a single vaccination, none of the vaccines completely prevented infection by SARS-CoV-2 Delta as evidenced by the detection of viral RNA in both the upper and lower respiratory tract of all animals (Figure 1C, D). However, on day 2 post challenge, sCPD9 vaccination diminished replicating virus to undetectable levels in the lungs; a feature none of the other groups exhibited (Figure 1E).

In principle, a booster dose 21 days after the initial vaccination improved vaccine efficacy based on virological evaluation following challenge infection (Figure 1F – H), even though these independent experiments cannot directly be cross-compared. Like the situation in the prime-only groups, viral RNA was detectable in all groups at all time-points and in both oropharyngeal swabs and lungs. However, all vaccine combinations significantly reduced virus RNA loads. Vaccination schedules involving sCPD9 yielded significantly lower viral RNA levels after challenge infection when compared to both two applications of mRNA or Ad-Spike vaccines (Figure 1F, G). Similarly, the levels of replication-competent virus in the lungs were significantly reduced in vaccinated animals on day 2 post challenge (dpc). Importantly, only sCPD9 booster vaccination reduced replicating virus levels below the detection, indicating sterilizing immunity following a booster vaccination with sCPD9 regardless of heterologous (mRNA) or homologous (sCPD9) priming (Figure 1H). On a virus transcript level, these results were confirmed by sequencing of bulk RNA extracted from the lungs (Figure S1B).

### LAV is superior in preventing inflammatory damage to the lung

To determine the degree of protection from virus-induced tissue damage and inflammation, infected hamster lungs were examined by histopathology. We found that after single vaccination, sCPD9 prevented inflammation and pneumonia more efficiently than other vaccines. sCPD9 vaccinated animals showed less consolidated lung areas (Figure 1M) and scores for lung inflammation, bronchitis and edema were clearly reduced (Figure 1I, J; S1C – F). Furthermore, sCPD9 vaccinated animals stood out for a marked presence of subepithelial plasma cells and lymphocytes which was not observed for other groups. Instead, animals that underwent other vaccination schedules displayed more prominent bronchial hyperplasia (Figure S2). A similar trend was observed for prime-boost regimens; however, particularly the mRNA vaccine benefited from the application of a second dose, improving the histological outcome compared to unvaccinated controls (Figure 1K, L; S1G – J). Nevertheless, application of two sCPD9 doses provided a superior outcome in all cases, as visualized by whole lung sections prepared at 5 dpc (Figure 1M; S2).

Concomitantly, bulk transcriptome analysis of hamster lungs from the prime-boost vaccinated animals showed a broad downregulation of infection- and inflammation-related genes in vaccinated compared to unvaccinated hamsters at both sampling time points (Figure S3). Consistent with the virological and pathological data, two sCPD9 vaccinations showed the strongest effect, followed by the heterologous mRNA+sCPD9 scheme. Likewise, in lung sections (5dpc) from the prime-boost vaccination experiment, inflammatory changes in animals that received sCPD9 were reduced to a minimum (Figure 2A).

**FIG. 2.**
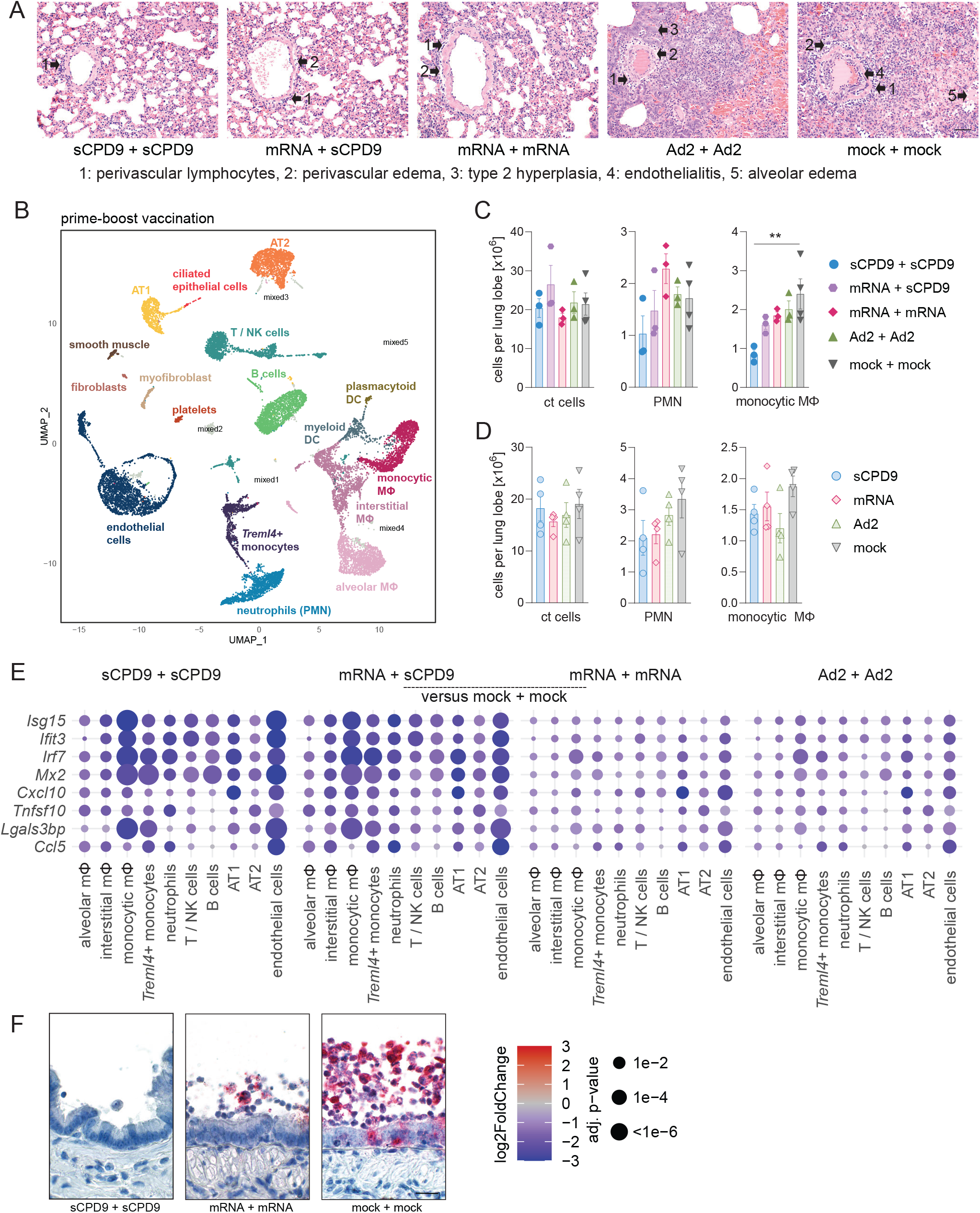
Pneumonia and pro-inflammatory transcriptional response is strongly reduced in vaccinated animals. **(A)** Haematoxylin-eosin-stained lung sections at 5 dpc from the prime-boost vaccination experiment identified perivascular lymphocytes (1), perivascular edema (2), metaplastic epithelial remodeling (3), endothelialitis (4), alveolar edema (5) as labeled by numbered arrows, groups as indicated. Scale bar = 90 μm. (**B**) Two-dimensional projections of single-cell transcriptomes using UMAP of lung cells (prime-boost experiment). Cells are colored by cell types as annotated based on known marker genes. **(C, D)** Manual cell count of isolated lung cells per lung lobe (ct cells), calculated numbers of indicated cell types (PMN, monocytic macrophages) based on scRNA-seq-determined cell frequencies for the prime-boost experiment **(C)** and prime experiment **(D)**. Bar plots with mean ± SEM, symbols representing individual hamsters (n = 3 – 4), ordinary one-way ANOVA and Tukey’s multiple comparisons test. ** p < 0.01. **(E)** Dotplots showing fold changes of gene expression in indicated cell types of the four prime-boost vaccination strategies compared to mock-mock vaccinated animals. Selected interferon-stimulated genes and pro-inflammatory cytokines are visualized as following. Coloration and point size indicate log2-transformed fold changes (FC) and p-values, respectively, in vaccinated compared to mock-mock vaccinated animals. Adjusted (adj) p-values were calculated by DEseq2 using Benjamini–Hochberg corrections of two-sided Wald test p-values. Genes are ordered by unsupervised clustering. **(F)** Localization of viral RNA by in situ-hybridization in a longitudinal section of a bronchus at 2 dpc. Red signals: viral RNA, blue: hemalaun counterstain. Note varying numbers of macrophages in lumen. Scale bar = 30 μm.

To gain a more detailed understanding regarding the effect of vaccination on lungs of SARS-CoV-2 infected hamsters, we performed single-cell RNA-sequencing (scRNA-seq) of lung samples from both vaccination experiments (Figure 2B; S4A). In sCPD9+sCPD9 vaccinated animals, pulmonary recruitment of monocytic macrophages was significantly reduced at 2 dpc. Pulmonary neutrophil recruitment, which is usually initiated within hours after pathogen contact (*31*), was also reduced, albeit to a lesser extent (Figure 2C; S4B). A similar effect, although less pronounced, was observed in the sCPD9 prime-only experiment (Figure 2D; S4C).

Single-cell RNA-sequencing allowed us to define which cell types exhibit the most prominent changes in gene activation in vaccinated animals upon virus challenge. Particularly, the interferon-stimulated genes (ISGs) known to be induced by SARS-CoV-2 infection (*32*) were broadly downregulated in vaccinated compared to unvaccinated animals, with monocytic macrophages, Treml4+ monocytes and endothelial cells appearing particularly responsive (Figure S5). Inflammatory mediators such as Cxcl10 or Tnfsf10 showed a more uniform pattern compared to ISGs (Figure 2E; S6A). Since macrophage subtypes showed markedly different gene expression patterns in the sCPD9+sCPD9 compared to the mRNA+mRNA and unvaccinated animals, we looked for the distribution of viral RNA within lungs (Figure 2F; S6B). In unvaccinated animals, the virus was generally detectable in large parts of the lungs, mRNA+mRNA vaccination reduced the occurrence to single patches, while in sCPD9+sCPD9 animals viral RNA was barely detectable by in-situ hybridization (Figure S6B). Notably, for the mRNA+mRNA regime, most detectable viral RNA appeared in macrophages (Figure 2F). Since virus uptake in macrophages causes their transcriptional activation (*32*), this can explain more abundant expression of pro-inflammatory genes in macrophages of mRNA+mRNA animals compared to those of sCPD9+sCPD9 vaccinated animals. Consistently, all effects were most pronounced after double sCPD9 vaccination, which could indicate that this strategy is also very successful in reducing the strong inflammatory responses that are responsible for both acute and long-term damage caused by SARS-CoV-2 infection.

### Humoral immunity against SARS-CoV-2 is most potent when the LAV is included in the vaccination regimen

To determine the potency of humoral responses, we quantified the ability of hamster sera collected before challenge (day 0) and on days 2 and 5 after challenge to neutralize the ancestral SARS-CoV-2 variant B.1. Following a single vaccination, the serum-neutralization capacity of sCPD9 vaccinees significantly exceeded all other groups at all time points (Figure 3A). Similarly, sera obtained from the sCPD9-vaccinated group provided superior neutralization of variants of concern B.1.351 (Beta), B.1.617.2 (Delta) and B.1.1.529 (Omicron, BA.1), especially at early time points when antibody titers are less influenced by challenge infection (Figure 3B, 3C). Particularly for Omicron BA.1, the neutralization capacity was considerably reduced in all groups (Figure 3D). Furthermore, sera from Ad2 spike-vaccinated hamsters had only weak neutralizing activity and sera from mRNA-vaccinated hamsters failed to neutralize SARS-CoV-2 prior to challenge infection (Figure 3D). This observation is consistent with studies showing that adenovirus vector vaccines can elicit stronger antibody responses than mRNA vaccines after a single dose administration (*33*–*35*). Generally, challenge infection resulted in an increase of neutralizing antibodies over time in all groups of hamsters (Figure 3A – D). However, regardless of the challenge effect, sCPD9 vaccinee sera neutralized all SARS-CoV-2 variants more efficiently at all time points after infection when compared to sera of otherwise vaccinated animals and significantly more efficiently than sera of unvaccinated hamsters (Figure 3A – D).

**FIG. 3.**
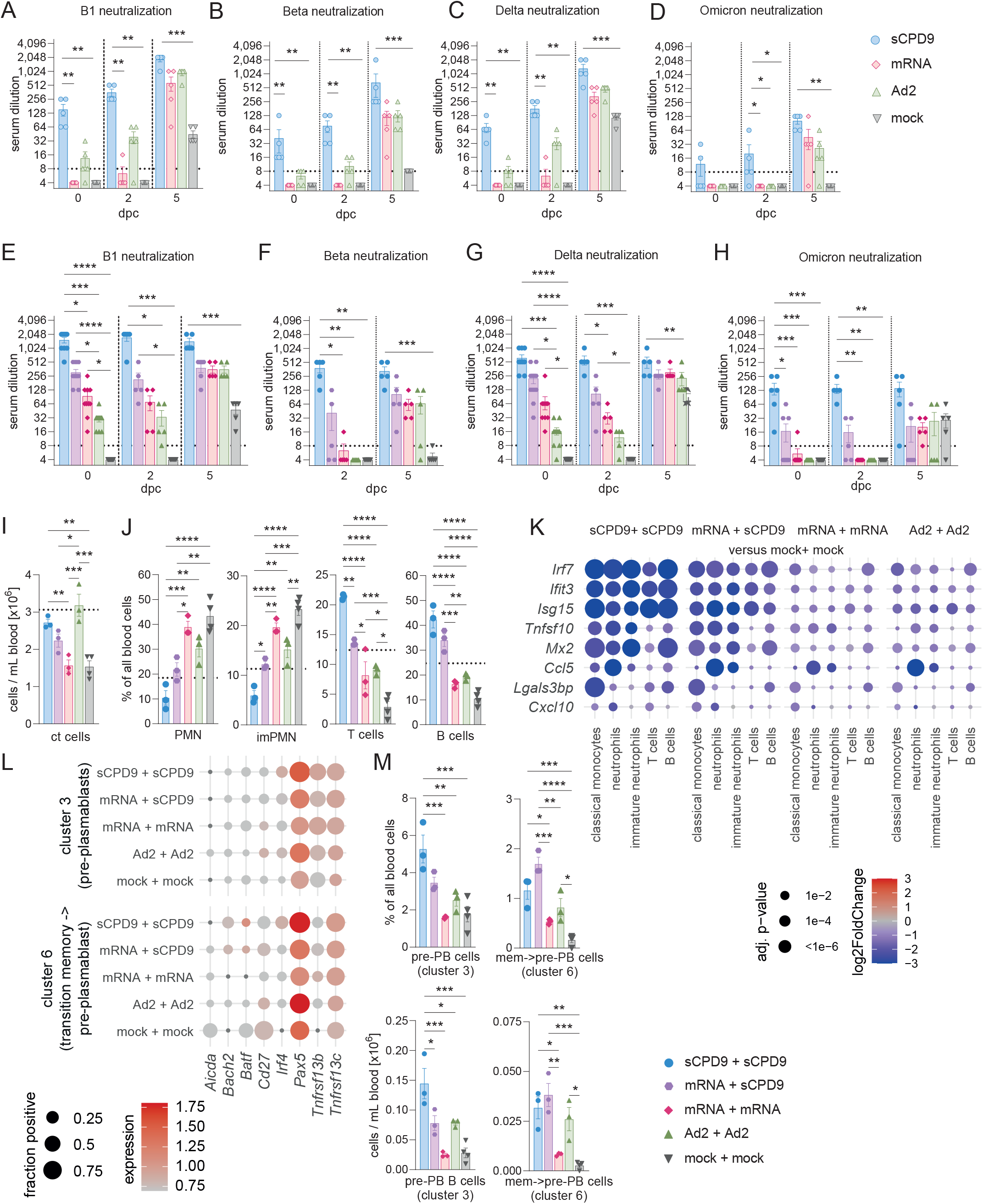
Antibody and cellular immune response in blood to vaccination and challenge. **(A – H)**, Serum neutralization titers of hamsters having received prime **(A – D)** and prime-boost **(E – H)** vaccination prior (0 dpc) and at 2- and 5-days post challenge with SARS-CoV-2. Neutralization capacity was tested against variant B1 **(A, E)**, Beta **(B, F)**, Delta **(C, G)** and Omicron **(D, H)**. The lower limit of detection was dilution 1:8 (dotted line) and the upper limit 1:2048. Bar plots with mean ± SEM, n = 5. Kruskal-Wallis and Dunn’s multiple comparisons test were performed. **(I – M)**, Analysis of cellular composition and gene expression by scRNA-seq at 2 dpc in blood of prime-boost vaccinated hamsters. Manual count of cells per mL blood **(I)**. Frequencies of indicated cell types among blood cells **(J)**. **(I, J)**, Dotted line marks the mean level found in naïve hamsters (n = 3, naïve hamster data derived and reprocessed from Nouailles et al., 2021). Bar plots with mean ± SEM, n = 3. One-way ANOVA and Tukey’s multiple comparison test were conducted. **(K)**, Dotplots showing fold changes of gene expression in indicated cell types of the four prime-boost vaccination strategies compared to mock-mock vaccinated animals. Selected interferon-stimulated genes and pro-inflammatory cytokines are visualized as following. Coloration and point size indicate log2-transformed fold changes (FC) and p-values, respectively, in vaccinated compared to mock-mock vaccinated animals. Adjusted (adj) p-values were calculated by DEseq2 using Benjamini–Hochberg corrections of two-sided Wald test p-values. Genes are ordered by unsupervised clustering. **(L)**, Dotplots showing expression of selected B cell development marker genes in the blood B cell subclusters shown in Fig. S9A. The size of the dot represents the fraction of cells in which at least one UMI of the respective was detected, the color is proportional to the average expression in those cells. **(M)**, Frequencies and numbers of pre-plasmablast (pre-PB) identified in B cell cluster 3 and memory to pre-plasmablast transitioning cells (mem->pre-PB) identified in B cell cluster 6. Bar plots with mean ± SEM, n = 3. One-way ANOVA and Tukey’s multiple comparison test were performed. * p < 0.05, ** p < 0.01,*** p < 0.001, **** p < 0.0001.

Hamsters that were boosted with sCPD9 or the mRNA vaccine produced measurably more neutralizing antibodies than those receiving prime-only vaccination. Contrarily, a boost vaccination with Ad2-spike was less efficient in enhancing antibody titers (Figure 3E – H). Overall, the booster vaccination increased serum neutralization capacity across different variants, with the strongest humoral response detected in animals that had received sCPD9 as booster (Figure 3E – H). Clearly, among the tested variants, Omicron BA.1 displayed the greatest ability to escape neutralization. It is however important to note that, contrary to the other vaccination regimens, prime-boost vaccination with sCPD9 provided hamsters with a significant ability to neutralize this SARS-CoV-2 variant (Figure 3H).

For a more detailed understanding of the differences between vaccination strategies, we evaluated cellular immune responses in blood of prime-boost vaccinated animals by scRNA-seq analysis. Unsupervised clustering followed by manual cell type annotation identified the expected major blood cell populations (Figure S7A). Manual blood cell counting identified significantly higher cell densities in sCPD9- and Ad2-vaccinated prime-boost groups relative to the controls, whilst heterologous mRNA-sCPD9 prime-boosted hamsters displayed a higher trend (Figure 3I). Combining these counts with the relative cell type composition obtained through scRNA-seq, we determined absolute cell numbers per mL of blood. Both relative and absolute cell numbers revealed substantial differences between vaccination strategies (Figure 3J; S7B – D). Frequencies of mature and immature neutrophils (imPMN), which are increased particularly in severe COVID-19 (*36*), but also in SARS-CoV-2-infected Syrian hamsters (*32*), were lowest for sCPD9+sCPD9-vaccinated animals (Figure 3J). Notably, neutrophil frequencies were also significantly lower for mRNA-sCPD9 and Ad2 vaccinated animals compared to unvaccinated hamsters. In contrast, the effectors of adaptive immunity, B, T and plasma cells, followed the opposite trend and displayed highest abundancies following the double sCPD9 regime (Figure 3J; S7B). Concordant with our observations in the lungs, genes related to infection and inflammation were broadly downregulated in myeloid cells of vaccinated animals relative to unvaccinated controls, particularly in neutrophils and monocytes (Figure 3K; S8).

To investigate activation of vaccine-induced immune memory, we first examined the single-cell transcriptome of circulating B cells, with a focus on B and plasma cells derived from the blood of prime-boost vaccinated hamsters; the subclustering of which displayed 8 populations (Figure S9A). Based on known marker genes (*37*, *38*), we first determined that cluster 8 represents plasmablasts/plasma cells, due to the presence of, for example, Prdm1 (which encodes Blimp-1) or Irf4. Based on intermediate Prdm1 levels, and the presence of Tnfrsf17 and Tnfrsf13b, cluster 3 was considered to be enriched for pre-plasmablasts (pre-PB). With higher levels of Pax5, Cd19, Cd27, Bach2 or Aicda, and lower levels of Prdm1, Xbp1 or Spib, we identified cluster 6 to primarily represent a population that transits from memory B cells towards pre-plasmablasts (mem->pre-PB) (Figure S9B). Since activation of pre-plasmablasts is prerequisite to potent memory recall responses, we investigated gene expression patterns in vaccinated versus unvaccinated animals in pre-PB (cluster 3) and mem->pre-PB (cluster 6) cells. Probing a set of genes involved in B cell regulation, we found upregulation of Irf4, Pax5 and Tnfrsf13b/c in pre-PB (cluster 3), while in mem->pre-PB (cluster 6) cells Bach2, Irf4, Pax5 and Tnfrsf13b/c were upregulated while Aicda, Batf and Cd27 were downregulated (Figure 3L; S9C). Although these differences do not necessarily directly reflect the process of B cell activation towards plasmablasts (e.g., Pax5 is downregulated in plasmablasts, but upregulated here), they could be interpreted as an early “memory recall gene expression signature” visible in the scRNA-seq data at 2 dpc. Notably, expression of this signature in blood B cells was strongest in hamsters that had received homologous or heterologous prime-post vaccination with sCPD9, which likewise generated highest antibody titers (Figure 3E – H). In line with these findings, the numbers and frequencies amongst blood cells of pre-PB (cluster 3) and mem->pre-PB (cluster 6) cells were significantly higher in sCPD9+sCPD9 vaccinated hamsters (Figure 3M).

### LAV enhances T cell proliferation in response to challenge with SARS-CoV-2

To investigate the occurrence of T cell memory recall responses, we proceeded with subclustering T and NK cells. To this end, we assayed CD4+, CD8+, and proliferating T cells in blood (Figure 4A, B; S10, S11). As expected, CD4+ and CD8+ T cells recapitulate the pattern generally observed for T cells (Figure 4A, 3J). Analyses of gene expression indicative of proliferation (Mki67, Top2a), naïve or central memory status (Sell, Ccr7, Lef1, Il7r) and activation of T cells (Cd69, Cd44, Klrg1, Tnfsf9, Icos, Cd40lg) revealed that most T cells in the blood displayed either naïve or central memory phenotypes (cluster 0 – 4, Figure S10B – E). In blood taken at 2 dpc, type 1-immunity effector genes (Tbx21, Gzma, Gzmb, Faslg, Ifn) were only expressed by NK cells (cluster 5, Figure S11). The proliferating T cell population consisted of activated T cells expressing memory markers, such as Il7r (cluster 6, Figure S11). Of note, the concentration and frequencies of proliferating T cells, albeit generally small, were significantly increased after heterologous vaccination, when compared to unvaccinated hamsters (Figure 4B). Analysis of gene expression in cluster 6 revealed that the fraction of cells expressing proliferation markers as well as the expression level of proliferation associated genes was higher when animals were vaccinated, and highest when sCPD9 was included in the vaccination regimen (Figure 4C).

**FIG. 4.**
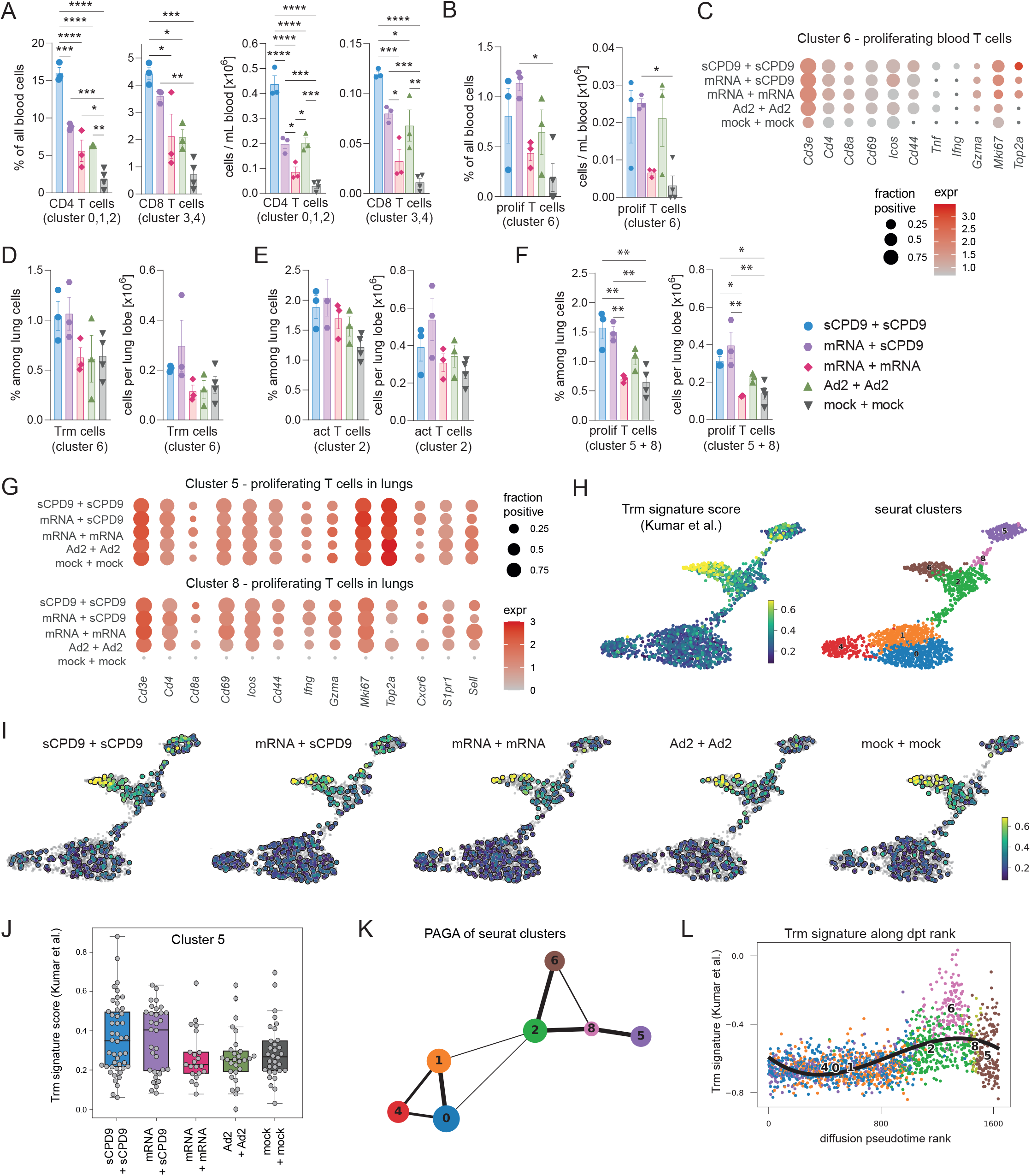
T cell responses in blood and lung in response to vaccination and challenge. **(A – K)**, Analysis of T cell subsets by scRNA-seq at 2 dpc in blood and lungs of prime-boost vaccinated hamsters. **(A)**, Frequencies and numbers of CD4 (cluster 0,1,2) and CD8 (cluster 3,4) T cells in blood. **(B)**, Frequencies and numbers of proliferating T cells (cluster 6) in blood. **(A, B)**, Bar graph with mean ± SEM, n = 3-4. Ordinary one-way ANOVA and Tukey’s multiple comparisons test. **(C)**, Dotplots showing expression of selected T cell marker genes in blood in cluster 6 derived from T and NK subcluster analysis in Fig. S10A. The size of the dot represents the fraction of cells in which at least one UMI of the respective was detected, the color is proportional to the average expression in those cells. **(D – F)**, Frequencies and numbers of (**D**) tissue-resident memory T cells (Trm, cluster 6), **(E)** activated T cells (Tact, cluster 2), **(F)** proliferating T cells (prolif T cells, cluster 5+8) in lungs. Ordinary one-way ANOVA and Tukey’s multiple comparisons test. Bar graph with mean ± SEM, n = 3-4. **(G)**, Dotplots showing expression of selected T cell marker genes in the lungs in cluster 5 and 8 derived from T and NK subcluster analysis in Fig. S12A. The size of the dot represents the fraction of cells in which at least one UMI of the respective was detected, the color is proportional to the average expression in those cells. **(H, I)**, Trm gene set (*Kumar et al., ref. 40*)enrichment in cells from selected T cell subclusters ranging from a low (purple) to a high (yellow) expression score over all groups (**H**) for individual groups as indicated (**I**). (**J**), Trm signature score in individual cells of cluster 5 in groups as indicated. Box, 25th to 75th percentile and whiskers Min to Max. (**K**) Partition-based graph abstraction (PAGA) where each node represents a cluster and edges to which extent two clusters are possibly connected. The size of the nodes corresponds to the number of cells in the cluster and the thickness of the lines is proportional to the connectivity. **(L)**, Trm signature (*Kumar et al., ref. 40*) score as a function of diffusion pseudotime rank with black line showing a polynomial fit of degree three.

Next, we examined whether different prime-boost vaccination strategies differed in the ability to re-activate tissue-resident memory T cells (Trm) in lungs of Syrian hamsters (*39*, *40*). To characterize pulmonary T cell subsets, we subclustered the initial T and NK cell clusters into 10 subclusters (Figure S12A). In analogy to the blood T cell readouts, we identified clusters 3, 7 and 9 as NK cells based on gene expression (Nkg7-true, Cd3-false), cluster 4 as CD8+ T cells (Cd3e, Cd8a), clusters 0, 1, 2, 6 as CD4+ T cells (Cd3e, Cd4) and clusters 8 and 5 as proliferating T cells (Cd3e, Mki67, Top2a) (Figure S12B, C). In line with NK cell gene expression pattern in the blood, NK cells located in the lungs likewise presented the most pronounced type 1-immunity effector gene expression (Tbx21, Faslg, Gzma, Gzmb) as well as Klrg1 (Figure S12D, E). Cluster 2 was dominated by CD4+ T cells and displayed a mixed phenotype of effector, activation and memory gene markers (Figure S12B – E, S13A, B). At 2 dpc of prime-boost vaccinated hamsters, most identified CD8+ T cells (cluster 4) were primarily of naïve or central memory type, whereas few CD8+ T cells scattered into the CD4+ T cell dominated clusters 5 and 8 of proliferating T cells. CD4+ T cells in clusters 0 and 1 were reminiscent of naïve or central memory type (Sell, Ccr7, Lef1, Il7r, Tcf7, S1pr1) (Figure S13A). In cluster 6, we did not find genes associated with naïve or central memory-associated signatures (Figure S13A) but combined and strong expression of T cell-homing and tissue retention genes (Cxcr6, Rgs1, Prdm1 (Blimp-1), Znf683 (Hobit), Itga1 (CD49a) and Itgae (CD103)), a signature indicative of Trm status (Figure S13B, C, S14). At 2 dpc, Trm (cluster 6), activated (cluster 2) and proliferating (cluster 5 and 8) T cell populations were small and represented less than 2 % of all lung cells in each case (Figure 4D – E). Trm (cluster 6) and activated T cells (cluster 2) trended towards higher frequencies and numbers in the lungs of sCPD9-vaccinated hamsters (Figure 4D, E). Gene expression analysis of Trm (cluster 6) showed that the gene expression level and cell fraction expressing Cxcr6, a prominent tissue homing receptor, was highest in sCPD9-vaccinated groups while lymph node retention receptor S1pr1 was least detected here (Figure S15A). Across activated T cells (cluster 2), gene expression of activation and effector genes was more uniform between groups and, thus, likely independent of prior vaccination (Figure S15A). Notably, like in blood, at 2 dpc numbers and frequencies of proliferating T cells were significantly higher in vaccinated groups and highest when animals had received sCPD9 as part of their vaccination regimen (Figure 4F). However, contrary to proliferating T cells in the blood, their lung counterparts expressed higher levels of effector genes such as Ifng and Gzma (Figure 4G). Moreover, when we scored the seurat clusters for a published human Trm gene set (*41*), we observed a subset of cells in cluster 5 (proliferating T cells) with a high Trm signature score (Figure 4H). At 2 dpc the Trm signature score in proliferating T cells (cluster 5) was remarkably higher when sCPD9 was part of the vaccination strategy (Figure 4I, J). In cluster 8 (proliferating T cells), overall cell numbers were too low to generate interpretable scores, and no cells from unvaccinated animals were identified (Figure S15B). To evaluate the extent to which the clusters are connected, we used a partition-based graph abstraction (PAGA) approach (*42*), which indicates particularly strong cellular connectivity between clusters 2, 8 and 5 and clusters 2 and 6 as well as a possible connection between cluster 6 and 8 (Figure 4K). Ordering cells according to global expression similarity by diffusion pseudotime (*43*) and plotting this rank against the Trm signature further corroborates a path between clusters 2/6 and clusters 8 and 5, which are accompanied by variable Trm-like gene expression (Figure 4L; S15C, D). Overall, these findings suggest that a subset of proliferating T cells is Trm recall-derived and activated in response to SARS-CoV-2 challenge infection in sCPD9-boosted hamsters.

### LAV induces superior mucosal immunity against SARS-CoV-2

In addition to potent T cell memory and humoral immunity, induction of protective mucosal immunity is a distinguishing property of LAVs that are administered locally at the natural site of virus replication (*44*). However, in case of SARS-CoV-2, induction of limited mucosal antibody responses following mRNA vaccination is reported (*45*, *46*). To assess induction of protective mucosal immune responses in the upper respiratory tract of hamsters that received different vaccine regimens, we measured SARS-CoV-2 spike-specific IgA levels after prime-only vaccinations. We found that sCPD9 vaccinated animals harbored considerably larger quantities of IgA in nasal washes prior challenge and at all tested time points post challenge (Figure 5A). Notably, challenge infection further boosted levels of SARS-CoV-2 spike-specific IgA antibodies in sCPD9-vaccinated animals and induced detectable quantities in mRNA- and Ad2-vaccinated animals, albeit with clear differences in favour of sCPD9 vaccinees at all time points. To confirm the protective effect of these antibodies, we performed microneutralization assays against authentic SARS-CoV-2 (variant B.1) with nasal washes obtained from the prime-boost experiment. Overall, the results obtained here recapitulated the IgA measurements. sCPD9-vaccinated animals exhibited markedly higher neutralization capacities at both 2 and 5 dpc (Figure 5B). Accordingly, we were able to identify IgA-positive lymphocytes in the nasal mucosa of vaccinated animals (Figure 5C). Protection at the mucosal level was also confirmed by histopathology of the nasal mucosa and SARS-CoV-2 localization by immunohistochemistry (Figure 5D, E). Histopathological scoring of nasal mucosa indicated that sCPD9-vaccinated animals displayed lesser affected tissue areas, lower damage levels and diminished immune cells recruitment (Figure 5D, E; S16A). Overall representing better protection from infection of the nasal mucosa with SARS-CoV-2. To further evaluate putative beneficial vaccination and mucosal immunity effects on the nasal cellular compartment, we resorted to scRNA-seq of nasal tissue cells. First, we annotated cell types in the nasal mucosa from the prime-only experiment based on previously published marker genes (Figure S16B, (*47*)). Differential gene expression analysis per cell type showed that particularly in nasal neuronal cells, interferon-stimulated genes such as Isg15, Oasl2 or Rsad2 were less expressed in vaccinated animals and particularly in sCPD9 vaccinees (Figure 5F; S17). This could be of particular importance for the efficiency of the vaccine, since bystander responses of neuronal cells in the olfactory epithelium are connected to loss smell upon SARS-CoV-2 infection (*48*). Taken together, we provide evidence that sCPD9, the live-attenuated vaccine employed here, provides superior protection against SARS-CoV-2 in both the lower and upper airways, making it a promising candidate for further investigation in clinical trials.

**FIG. 5.**
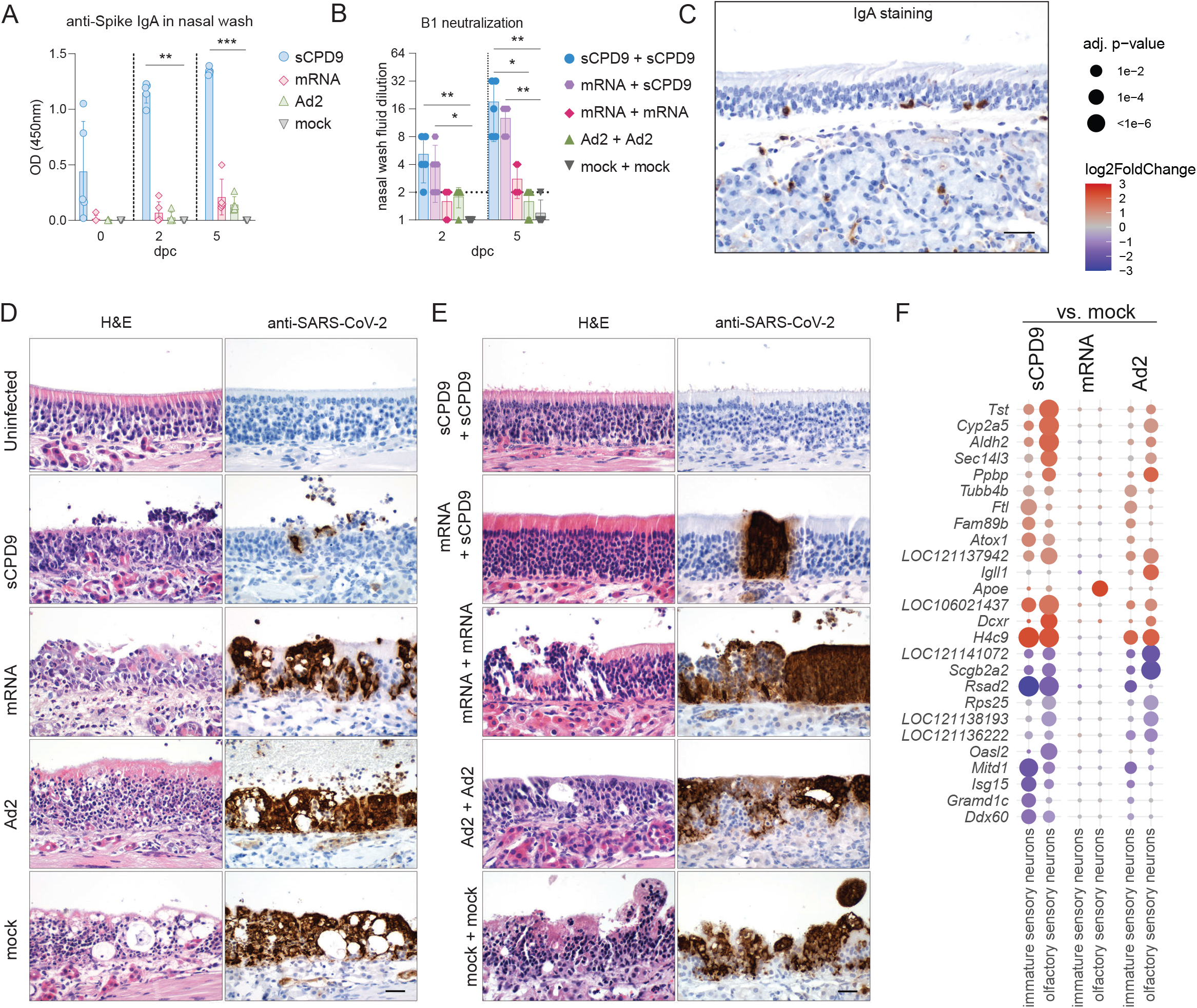
Protective effects on the mucosa and development of local immunity after vaccination. **(A)** ELISA detecting anti-spike IgA levels in nasal washes of prime vaccinated hamsters at indicated time points post challenge (dpc). Results display optical density reads at OD450nm. **(B)** Neutralizing capacity against B.1 of nasal wash fluids from prime-boost vaccinated hamster at indicated time points. (A – B) Bar plots show mean ± SD. Kruskal-Wallis and Dunn’s multiple comparisons test per time point. * p < 0.05, **p < 0.01, ***p < 0.001, ****p < 0.0001. **(C)**, Immunohistochemical staining of IgA in the olfactory epithelium and submucosal glands at 2 dpc. Scale bar = 60 μm. (**D**), shown are longitudinal histopathological sections of olfactory epithelium, with haematoxylin and eosin staining (left) and SARS-CoV-2 N protein immunohistochemistry (right) of the prime-only experiment at 2 dpc, with an additional section of an uninfected tissue. (**E**), as in (**D**) for the prime-boost vaccination experiment. **(E)**, as in **(D)** for the prime-boost vaccination experiment. **(F)**, Dotplots showing fold changes of gene expression in indicated cell types of the three prime vaccination strategies compared to mock vaccinated animals. Selected interferon-stimulated genes and pro-inflammatory cytokines are visualized as following. Coloration and point size indicate log2-transformed fold changes (FC) and p-values, respectively, in vaccinated compared to mock vaccinated animals. Adjusted (adj) p-values were calculated by DEseq2 using Benjamini–Hochberg corrections of two-sided Wald test p-values. Genes are ordered by unsupervised clustering.

## Discussion

The ongoing spread of SARS-CoV-2 variants is not efficiently controlled by current vaccination strategies. Although current vaccines, specifically mRNA vaccines, are highly effective in preventing severe and fatal disease, infection with newly emerging variants is not prevented and typically leads to symptomatic breakthrough infection with considerable virus loads even in triple-vaccinated individuals (*49*). To better control virus transmission and limit symptomatic infection, mucosal immunity at the site of virus entry is thought to be of paramount importance (*50*–*53*).

For the first time, our study presents a pre-clinical cross-platform vaccine comparison that includes a live attenuated vaccine. We find that specifically this live attenuated vaccine elicits superior protection from SARS-CoV-2 infection especially at mucosal sites of virus entry. This stands in agreement with previous preclinical COVID-19 vaccine studies using intranasal administration of LAV, protein-based or virus-vectored spike vaccines showing high efficacy in animal models (*29*) and induction of mucosal immunity (*54*–*60*). Our observations on improved immunity induced by heterologous immunization with an intramuscular mRNA vaccine followed by an intranasal sCPD9 boost are in line with recent studies that combine systemic priming followed by an intranasal boost with adenovirus vector vaccines (*61*, *62*). Importantly, virus-neutralizing anti-SARS-CoV-2 IgA at the nasal mucosa of vaccinated animals were found in considerably higher quantities in animals that had received the live attenuated vaccine candidate sCPD9. The great importance of local and broad antigenic IgA responses in acute viral infection and vaccination is known from previous work on other respiratory pathogens such as influenza A. Mucosal IgA exerts various functions in the local immune response, such as preventing release from mucins, blocking viral entry, preventing intracellular fusion of virus and endosomal membranes as well as inhibiting release of viruses from host cells (*63*). In line with this, overall protection from virus replication, tissue damage and lung inflammation were significantly better in sCPD9-vaccinated animals compared to all other groups. It is likely that these observations are a result of important hallmark features of live attenuated vaccines that include administration via the natural route of infection, presentation of the full antigenetic repertoire of the virus, and replication mimicking the target pathogen. Moreover, our scRNA-seq analysis of samples from blood, lungs and nasal mucosa of vaccinated and SARS-CoV-2 challenge-infected hamsters allowed us to gain further insights into the immunological consequences of different vaccine concepts. Across all important parameters, effects were strongest for sCPD9 vaccination in a prime-only setting. Similarly, in a prime-boost setting, double sCPD9 vaccination was superior to mRNA-sCPD9 vaccination, followed by double mRNA vaccination and double adenovirus vaccination. On the one hand, sCPD9-vaccinated animals showed substantially reduced induction of pro-inflammatory gene expression programs, which are a main cause for pathogenesis of COVID-19 (*64*). This was specifically true for cells of the innate immune system such as monocytes and macrophages, which typically display strong pro-inflammatory transcriptional responses upon SARS-CoV-2 uptake (*32*). If translatable to humans, this could mean a much higher chance for a mild or asymptomatic course of disease even in the case of infection with heterologous SARS-CoV-2 variants. Further, we provide evidence for reduced transcriptional activation of neuronal cells in the nasal mucosa. A recently published study on the infection of the olfactory epithelium in hamsters and humans showed that olfactory neurons do not get infected, but their reaction to infection of the neighbouring sustentacular cells would be connected to the well-described loss of smell upon SARS-CoV-2 infection (*48*).

Additionally, we detected several gene expression signatures that can be connected to activation of adaptive immune memory. We recorded enhanced development towards pre-plasmablasts derived from memory B cells and enhanced T cell proliferation in the blood of challenged animals by scRNA-seq. This points towards rapid activation of memory cells (*65*, *66*). Furthermore, analysis of local pulmonary immune responses by sc-RNA-seq allowed us to detect significantly increased numbers of proliferating T cells in lungs of hamsters that received sCPD9 as part of their vaccination regimen. A subset of those proliferating T cells shared a Trm-specific signature and showed connectivity to the identified Trm cluster. One possible explanation for the observation is that SARS-CoV-2-specific tissue-resident memory T cell seeding improves following sCPD9 vaccination and enables faster local recall responses, characterized by enhanced proliferating T cells in corresponding vaccine groups. However, in hamsters, antigen-specificity analysis of T cell receptors (TCRs) is currently not feasible and therefore ultimate proof is elusive at present. Nonetheless, LAV mimic natural infection, which is known to induce SARS-CoV-2 specific CD4^+^ Th1 cells secreting IFNγ, an antiviral effector cytokine (*67*). We detected IFNγ to be upregulated in proliferating pulmonary T cells, indicating SARS-CoV-2 challenge triggered development towards a TH1 effector cell type. While mucosal IgA induction remains most important in limiting infection and thus transmission, airway memory CD4^+^ T cells have been shown to significantly contribute to protection against other coronaviruses (*40*) and carry the potential to enhance the antigenic repertoire of any given mucosal vaccine against SARS-CoV-2. Similarly, earlier studies utilizing ovalbumin antigens and a combination of different vaccination routes indicated that not just IgA, but also general TH1-mediated immunity is enhanced upon mucosal delivery (*68*).

Our single-cell RNA-sequencing analysis has several limitations. This technique, as employed here, cannot fully capture processes such as reactivation of memory cells due to lack of surface markers and cell type-specific enrichment. Due to incomplete annotation of the Syrian hamster genome, we were not able to identify IgA-positive cells. Data quality of nasal mucosa cells was comparatively low due to the difficult dissociation of the tissue, which limits our observations at the site of initial infection.

An important and frequently discussed issue with live attenuated vaccines is their potential susceptibility to previously established immunity (*69*), which would restrict vaccine virus replication and potentially limit their use as booster vaccines after initial immunization by vaccination or natural infection. Our results indicate however, that sCPD9 does effectively boost immune responses and greatly improves protection when applied three weeks after initial mRNA vaccination. Importantly, sCPD9 enhances humoral immune responses, especially against known immune escape variants such as Beta and Omicron BA.1, while also improving the virological outcome of a heterologous challenge infection when applied as a booster three weeks after initial vaccination. This indicates a wide scope for the use of live attenuated vaccines in populations that exhibit an already high degree of baseline immunity induced by previous vaccination or infection; a situation clearly present in many parts of the world. Interestingly, in contrast to the single vaccinations, neutralization titers did not increase after challenge following dual vaccination with sCPD9. This result may indicate that challenge virus replication shortly after sCPD9 boost is suppressed to an extent that prevents further B-cell activation.

While our data shows superiority, and therefore promise for further development and refinement, of live attenuate vaccines, there is a caveat for extrapolating the results of preclinical animal trials to the situation in humans. Clearly, clinical studies regarding safety and efficacy of live attenuated vaccines are mandated to realistically assess the potential of these vaccines to combat the yet ongoing pandemic. Our work presented here can also guide some aspects in the evaluation of such clinical trials. Due to its high safety profile, sCPD9 was recently downgraded from biosafety level (BSL) 3 to BSL 2 by the relevant German state authority (Landesamt für Gesundheit und Soziales Berlin). This is an important step towards clinical application of a SARS-CoV-2 LAV as it will facilitate production of a clinical grade vaccine and greatly ease clinical trials in humans.

## Acknowledgments

We thank V.M. Corman, Charité, for his help in study design and prolific discussions on results and conclusions and S. Reiche, Friedrich-Loeffler-Institut, for providing anti–SARS-CoV-2 nucleocapsid antibody. We thank C. Thöne-Reineke for support in animal welfare and husbandry. GN thanks E. Nouailles for support in childcare. We thank the European Virus Archive, D. Bourquain from the Robert-Koch-Institut and C. Reusken from the National Institute for Public Health and the Environment for providing SARS-CoV-2 variants used in this study. VeroE6-TMPRSS cells were provided by the NIBSC Research Reagent Repository, UK. With thanks to Dr. Makoto Takeda. Figure 1A was created with BioRender.com.

## Author contributions

Conceptualization: DK and JT

Methodology: GN, JMA, PP, SP, GTA, MB, JB, AV, AL, FP, JK, CG, DN, HW, SDP, EW, DK, JT

Investigation: GN, JMA, PP, SP, GTA, JB, AV, AL, SS, NX, CL, RMV, AA, SH, HW, ADG, SDP, EW, DK, JT

Visualization: GN, JMA, PP, SP, GTA, JB, AV, AL, SDP, EW, DK, JT

Resources: GN, GC, ADG, CG, MW, DN, CD, JT

Funding acquisition: NO and JT

Project administration: JT

Supervision: GN, GC, CD, CG, ML, NB, MW, ADG, SDP, NO, EW, JT

Writing – original draft: GN, JMA, EW, DK, JT

Writing – review & editing: all

## Data and code availability

Raw sequencing data is available on GEO (https://www.ncbi.nlm.nih.gov/geo/query/acc.cgi?acc=GSE200596), along with bulk RNA-seq read count tables, and h5 matrices and Seurat objects for the single-cell RNA-seq data. Code is available through github at https://github.com/Berlin-Hamster-Single-Cell-Consortium/Live-attenuated-vaccine-strategy-confers-superior-mucosal-and-systemic-immunity-to-SARS-CoV-2.

