## Supplementary Figures for "A live attenuated vaccine confers superior mucosal and systemic immunity to SARS-CoV-2 variants"

**Fig. S1.**

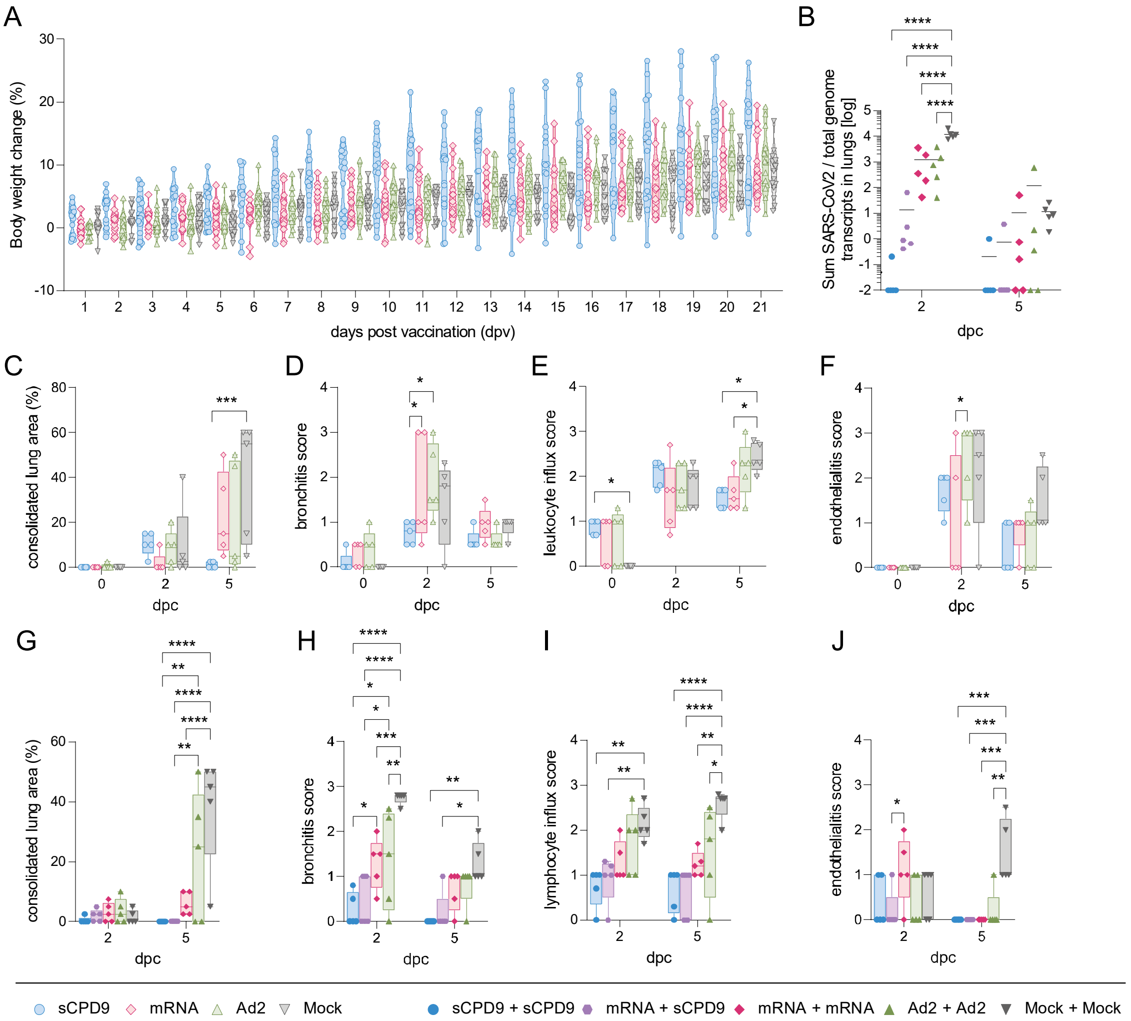


(**A**) Body weight development in percent of prime vaccinated hamsters were measured for 21 days until virus challenge time point and displayed according to vaccination group. Violine plot (truncated) with quartiles and median. (**B**) Prime-boost experiment: Relative expression of SARS-CoV-2 total canonical junction-spanning viral mRNA transcripts, compared to the total genomic transcripts, generated after bulk RNA sequencing analysis from homogenized lung tissue. Values are shown in log10 scale for both time-points analyzed. Scatter dot plot with mean. Two-way ANOVA (analysis of variance) and Tukey`s multiple comparison test. (**C** – **J**) Semi-quantitative scoring of histopathological findings of Syrian hamsters included in prime (**C** – **F**) and prime-boost setting (**G** – **J**): (**C**, **G**) Consolidated lung area found in the left lung lobe is displayed in percentage. (**D**, **H**) Bronchitis score accounts for severity of bronchial epithelial necrosis and bronchitis. (**E**, **I**) To consider local cellular immune response, pulmonary infiltration of neutrophils, lymphocytes and macrophages was assessed in the leukocyte influx score. (**F**, **J**) The extent of endothelialitis in the left lung lobe is indicated in the endothelialitis score. (**C** – **J**) Results are shown as box (25$^{th}$ to 75$^{th}$ percentile) and whiskers (Min to Max.) plots. Two-way ANOVA and Tukey`s multiple comparison test. $*p < 0.05$, $**p < 0.01$, $***p < 0.001$, and $****p < 0.0001$.

**Fig. S2.**

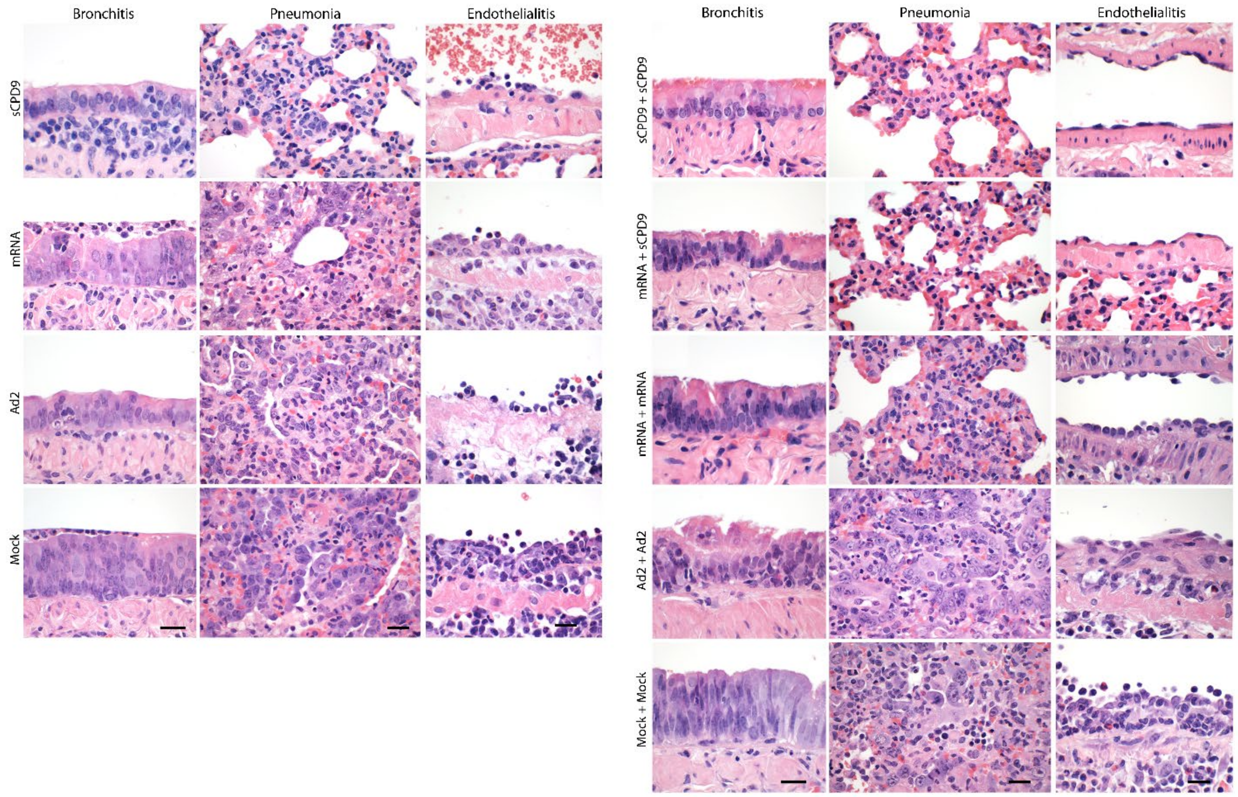


Representative histopathology of haematoxylin and eosin-stained lung sections from prime-only and prime-boost vaccination experiments at 5 dpc. For both experiments, columns show, from left to right, bronchitis, pneumonia affecting the respiratory alveoli and blood vessels with endothelialitis. In the prime-only approach, and in contrast to all other groups that developed necrosupperative and hyperplastic bronchitis, only sCPD9 vaccinated hamsters had negligible bronchitis in the presence of BALT-like subepithelial infiltration with lymphocytes and plasma cells. In the lungs, alveoli of sCPD9 vaccinated animals presented with much less consolidated respiratory parenchyma, with less infiltrating macrophages and neutrophils. Only Ad2 and mock-vaccinated animals developed marked alveolar metaplastic remodeling, indicating regeneration after necrosis of alveolar epithelia. Endothelialitis was milder in all vaccinated groups compared to mock-vaccinated animals. In the prime-boost experiments, hyperplastic bronchitis was mildest in sCPD9+sCPD9 and mRNA+sCPD9 vaccinated hamsters. Consolidation of respiratory parenchyma and alveolitis were least severe in both sCPD9 boostered groups. Metaplastic epithelial remodeling was particularly pronounced in Ad2-Ad2 vaccinated animals. Endothelialitis was strongly reduced to similar degrees in all boostered groups. Scale bars = 20 µm for each column.

**Fig. S3.**

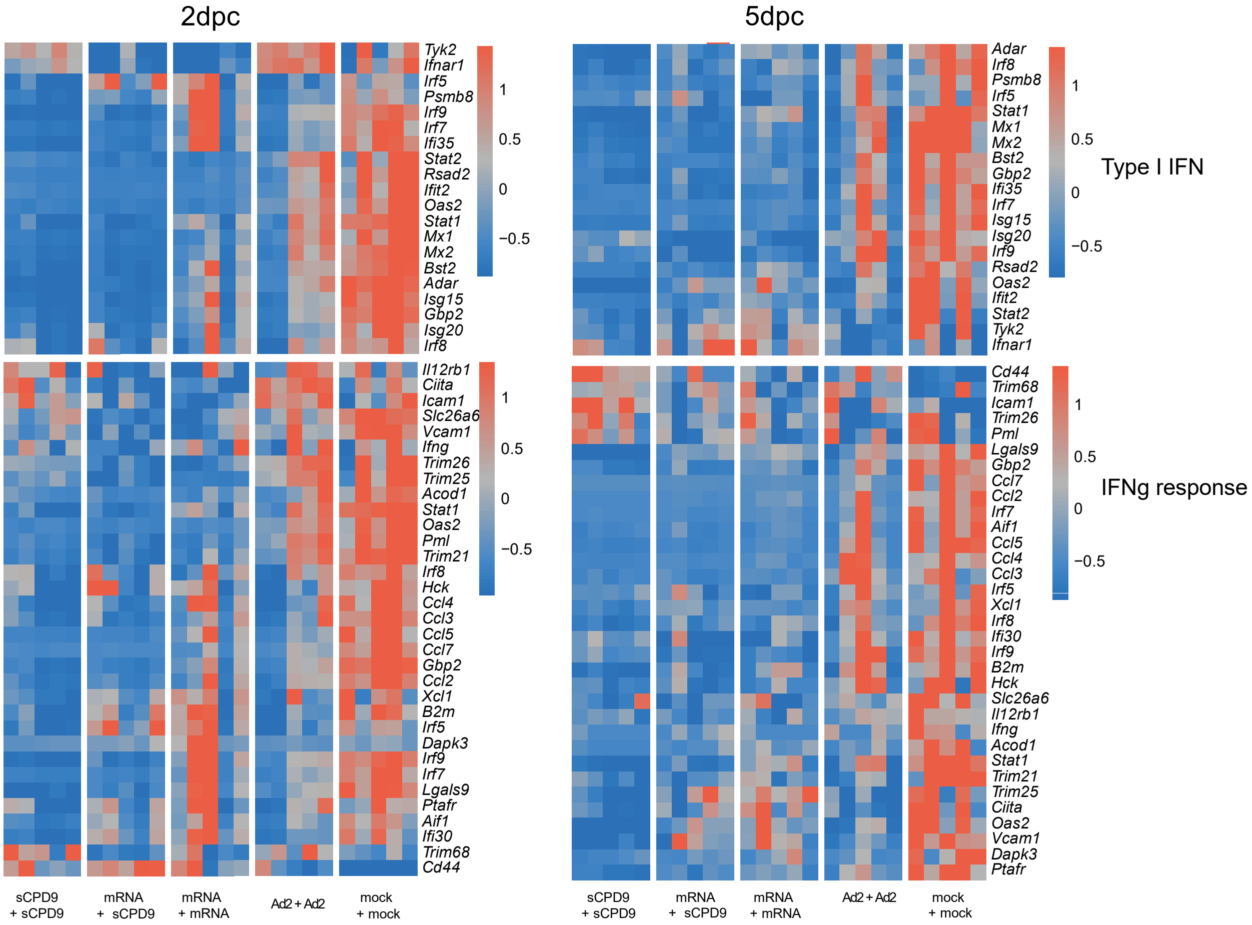


Heatmaps of differently expressed genes involved in the immune response at 2- and 5-days post challenge (dpc) with SARS-CoV-2 were analyzed in the lung tissue of Syrian hamsters after 4 different vaccination strategies using bulk RNA-sequencing. Genes involved in the type I IFN signaling and related to cellular response to IFNγ were as previously described (*78*). Columns represent samples and rows genes. Shown are z-scores of DESeq2-normalized data and color scale ranges from red (10 % upper quantile) to blue (10 % lower quantile), showing up- or down-regulation in expression of the selected genes. N = 5 animals per group.

**Fig. S4.**

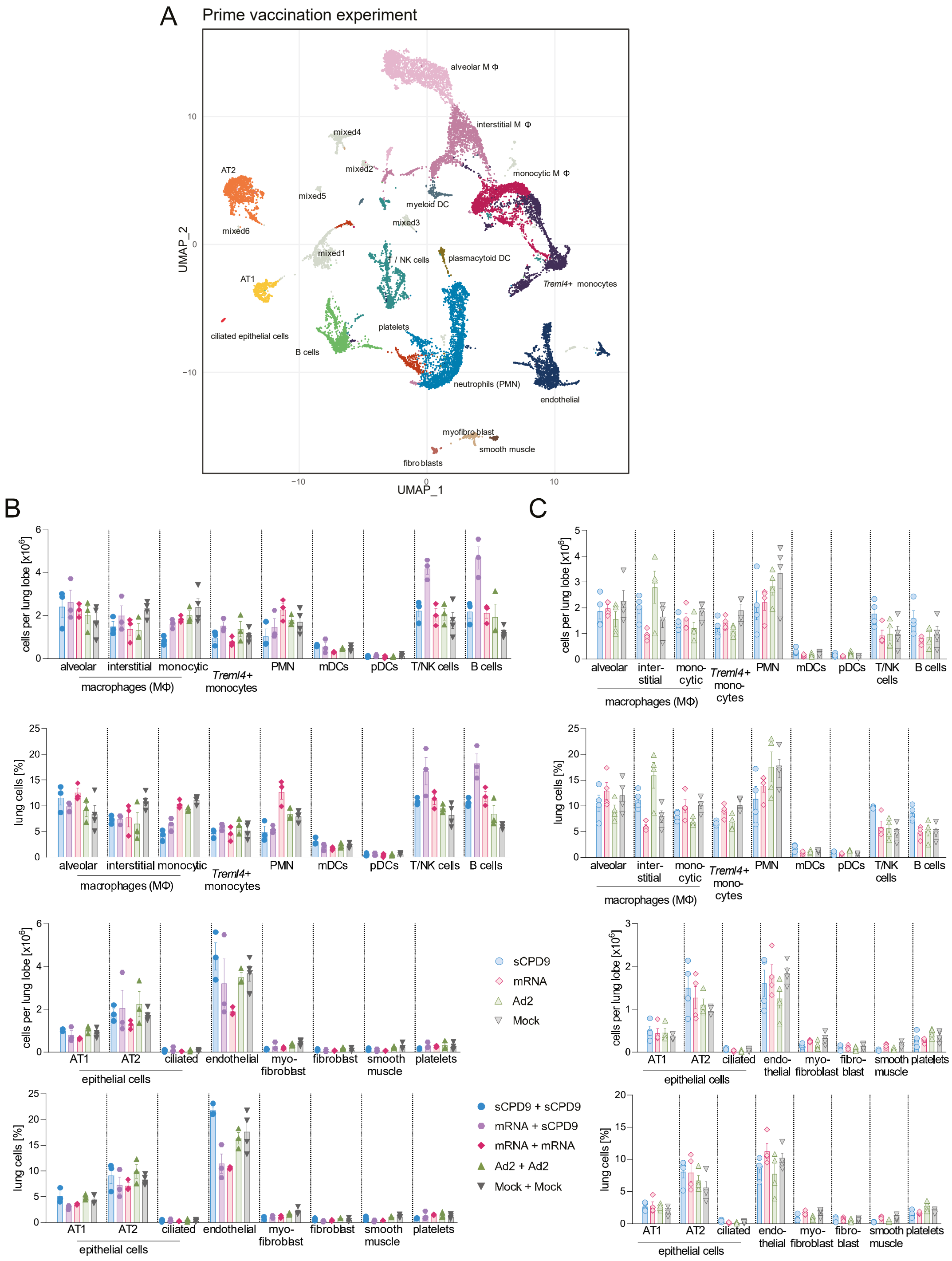


(**A**) Two-dimensional projections of single-cell transcriptomes using UMAP of lung cells from the prime vaccination experiment. Cells are colored by cell types as annotated based on known marker genes. (**B** – **C**) Numbers and percentage of cellular components per lung lobe for (**B**) prime-boost experiment and (**C**) prime-only vaccination experiment. Numbers and frequencies of PMN and monocytic MΦ from Fig. S4B are also shown in Fig. 2C and from Fig. S4C in Fig. 2D.

### Fig. S5.

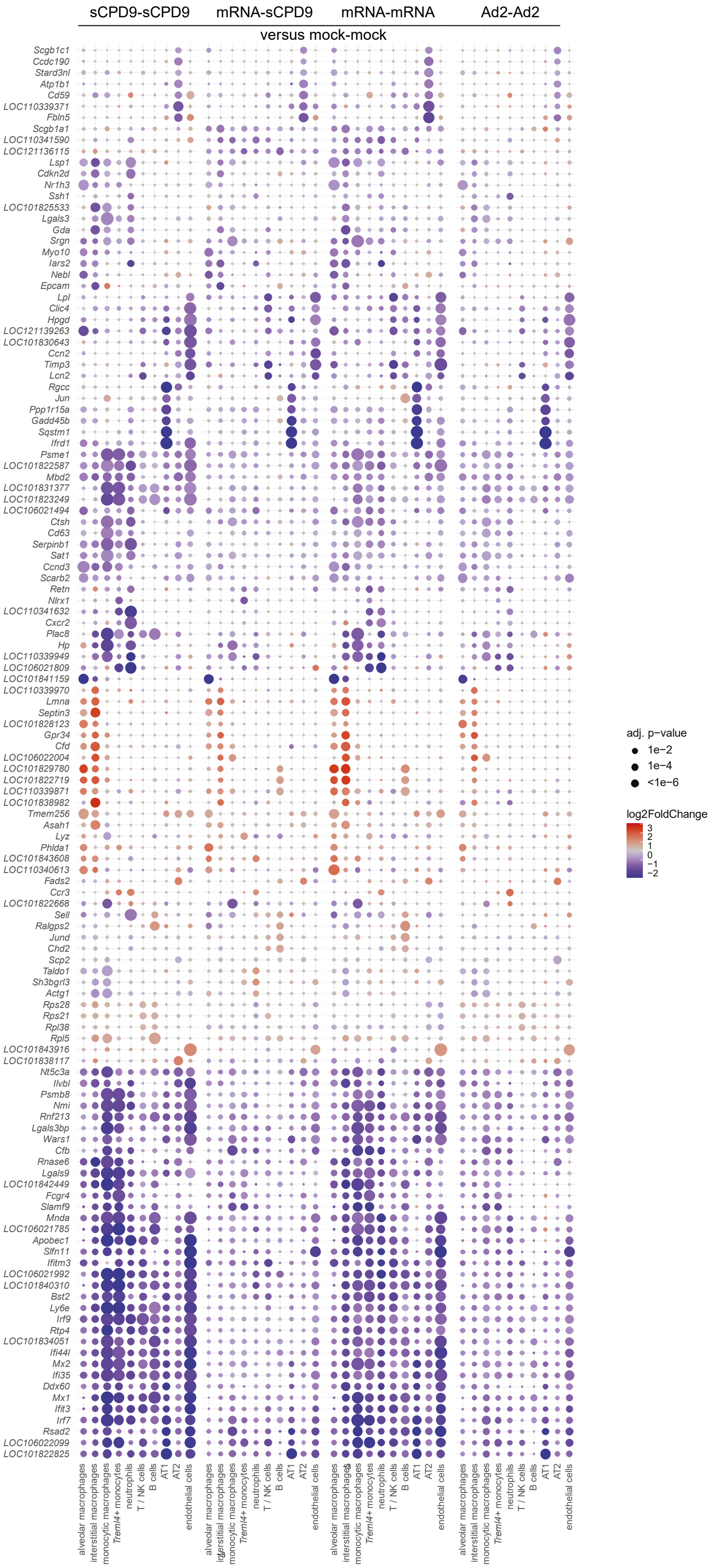


Dotplot showing fold changes of gene expression in indicated cell types of prime-boost vaccinated compared to mock-mock vaccinated animals. Coloration and point size indicate log2-transformed fold changes (FC) and p-values, respectively, in vaccinated compared to mock-mock vaccinated animals. Adjusted (adj) p-values were calculated by DEseq2 using Benjamini–Hochberg corrections of two-sided Wald test p-values. Genes are ordered by unsupervised clustering.

**Fig. S6.**

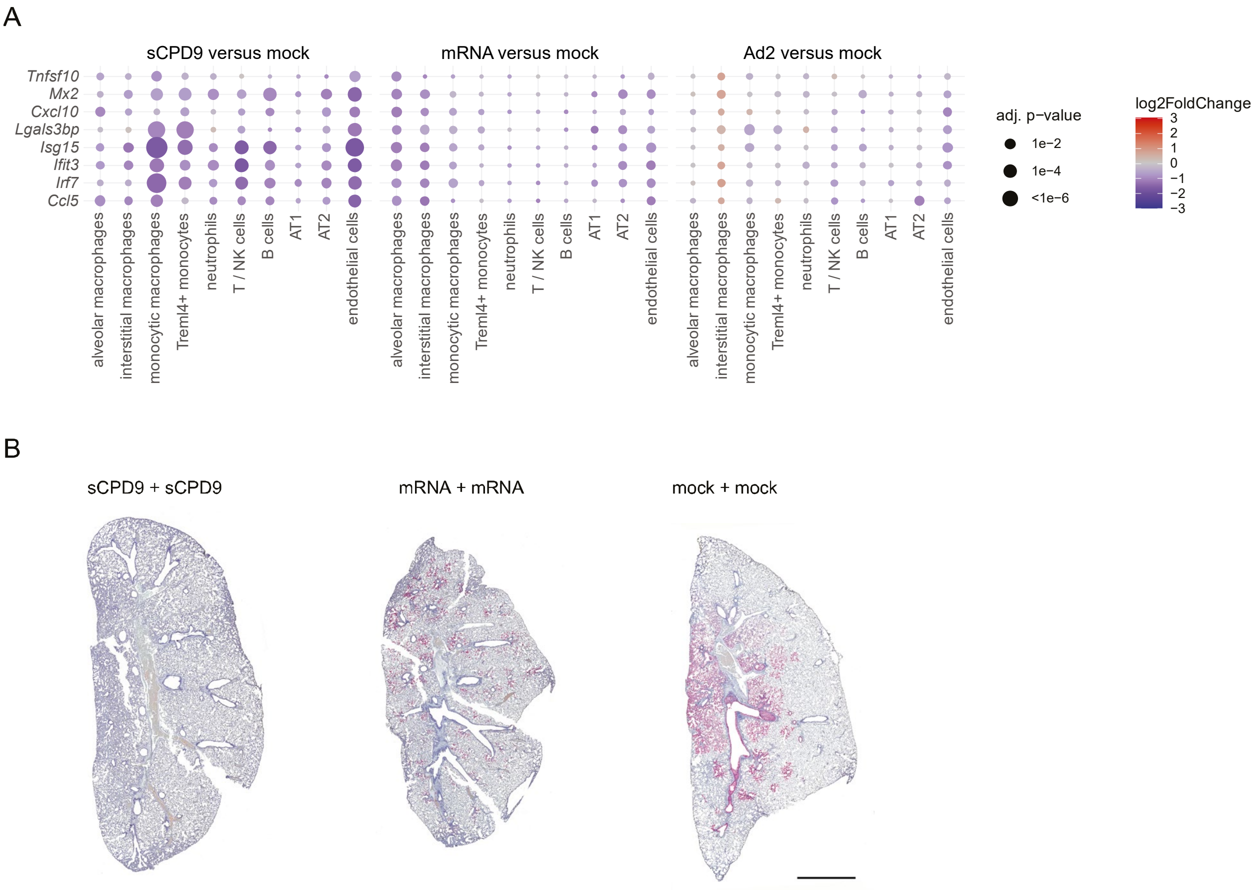


(**A**) Dotplots showing fold changes of gene expression in indicated cell types of the three prime vaccinations compared to mock vaccinated animals. Selected interferon-stimulated genes and pro-inflammatory cytokines are visualized as following. Coloration and point size indicate log2-transformed fold changes (FC) and p-values, respectively, in vaccinated compared to mock-mock vaccinated animals. Adjusted (adj) p-values were calculated by DEseq2 using Benjamini–Hochberg corrections of two-sided Wald test p-values. Genes are ordered by unsupervised clustering. (**B**) Localization of viral RNA (ribonucleic acid) by in situ-hybridization in scanned sections of left lung lobes at 2 dpc in indicated vaccination groups. Red signals: viral RNA, blue: hemalum counterstain. Scale bar represent 3 mm.

**Fig. S7.**

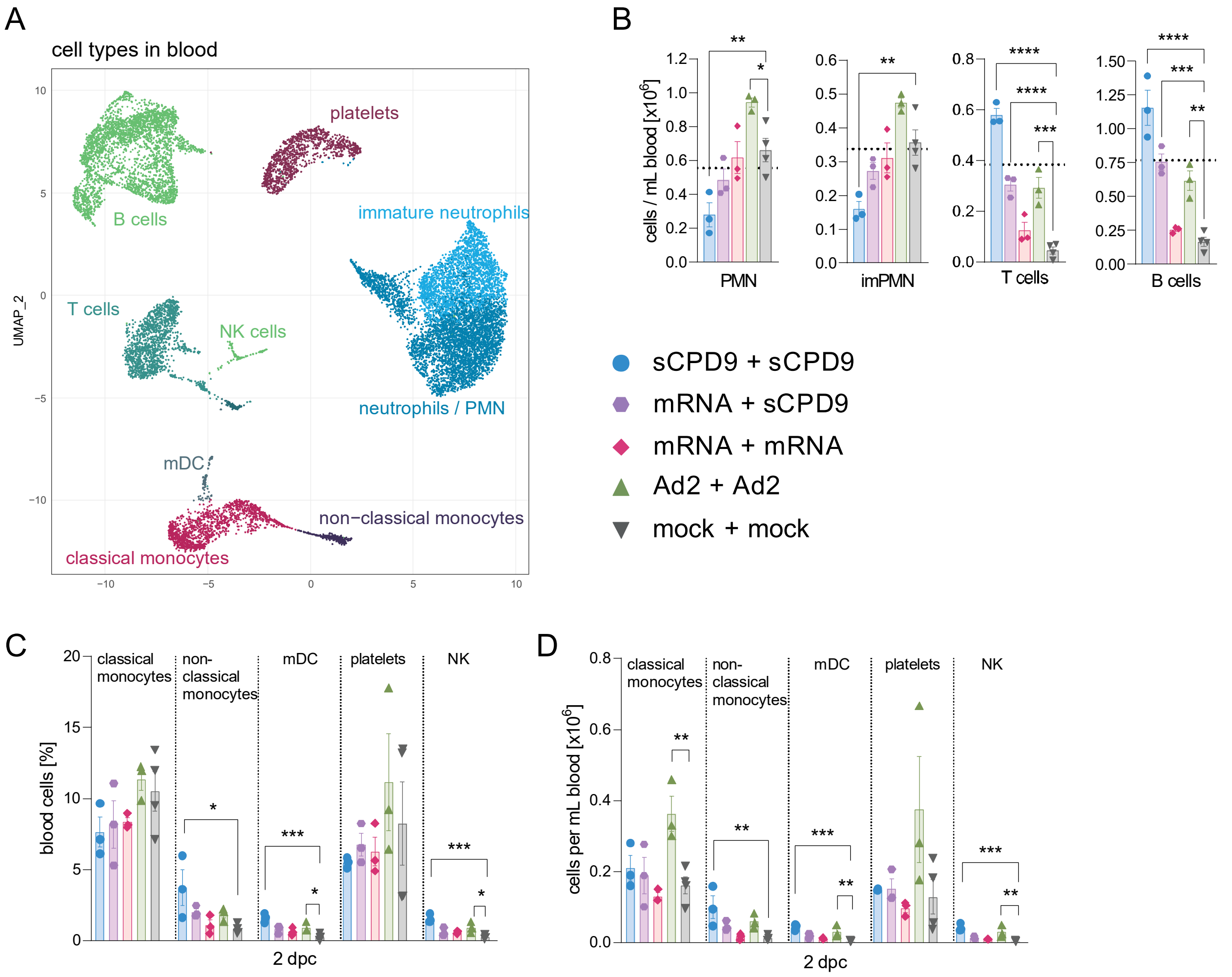


(**A**) Two-dimensional projections of single-cell transcriptomes using UMAP of blood cells from prime-boost experiment. Cells are colored by cell types as annotated based on known marker genes. (**B**, **D**) Numbers of cellular components per ml blood for the prime-boost experiment. (**C**) Percentage of blood cellular components for the prime-boost experiment. (**B** – **D**) One-way ANOVA and Dunnett's multiple comparisons test against mock-mock group per cell type are shown. $*p < 0.05$, $**p < 0.01$, and $***p < 0.001$.

#### Fig. S8.

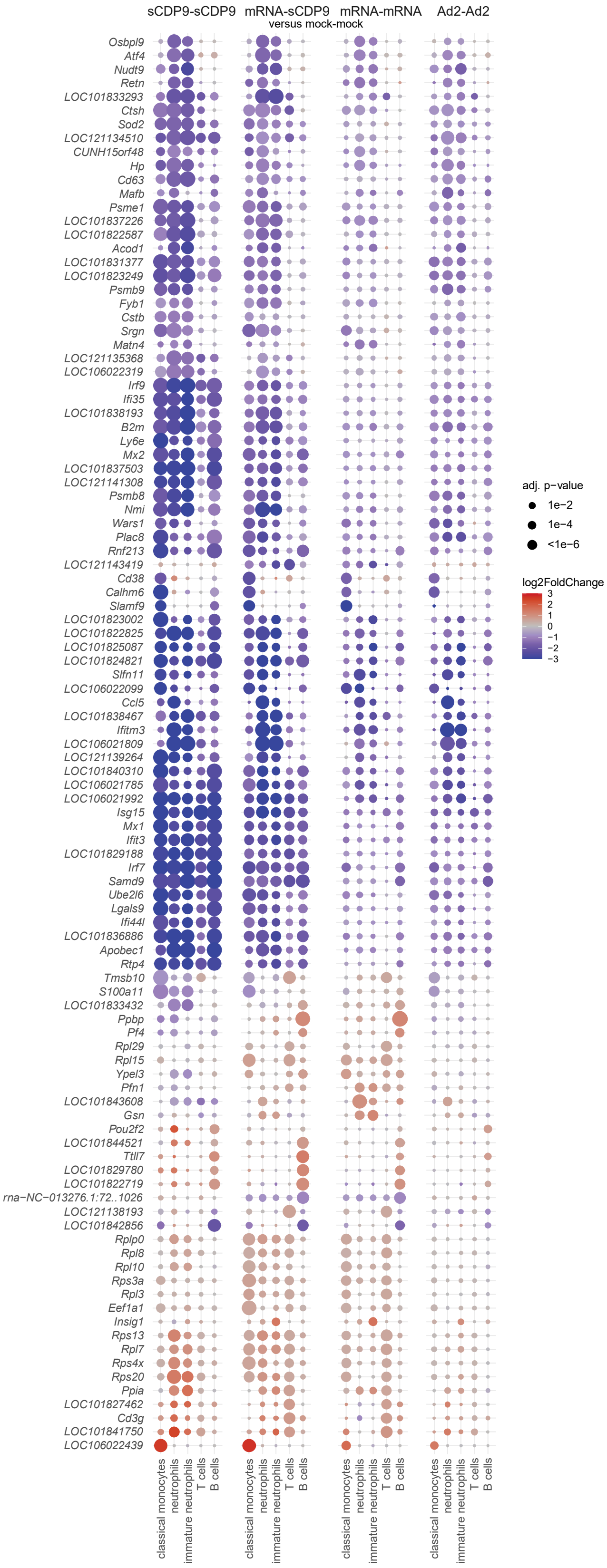


Dotplots showing fold changes of gene expression in indicated cell types of prime-boost vaccinated compared to mock-mock vaccinated animals. Coloration and point size indicate log2-transformed fold changes (FC) and p-values, respectively, in vaccinated compared to mock-mock vaccinated animals. Adjusted (adj) p-values were calculated by DEseq2 using Benjamini–Hochberg corrections of two-sided Wald test p-values. Genes are ordered by unsupervised clustering.

**Fig. S9.**

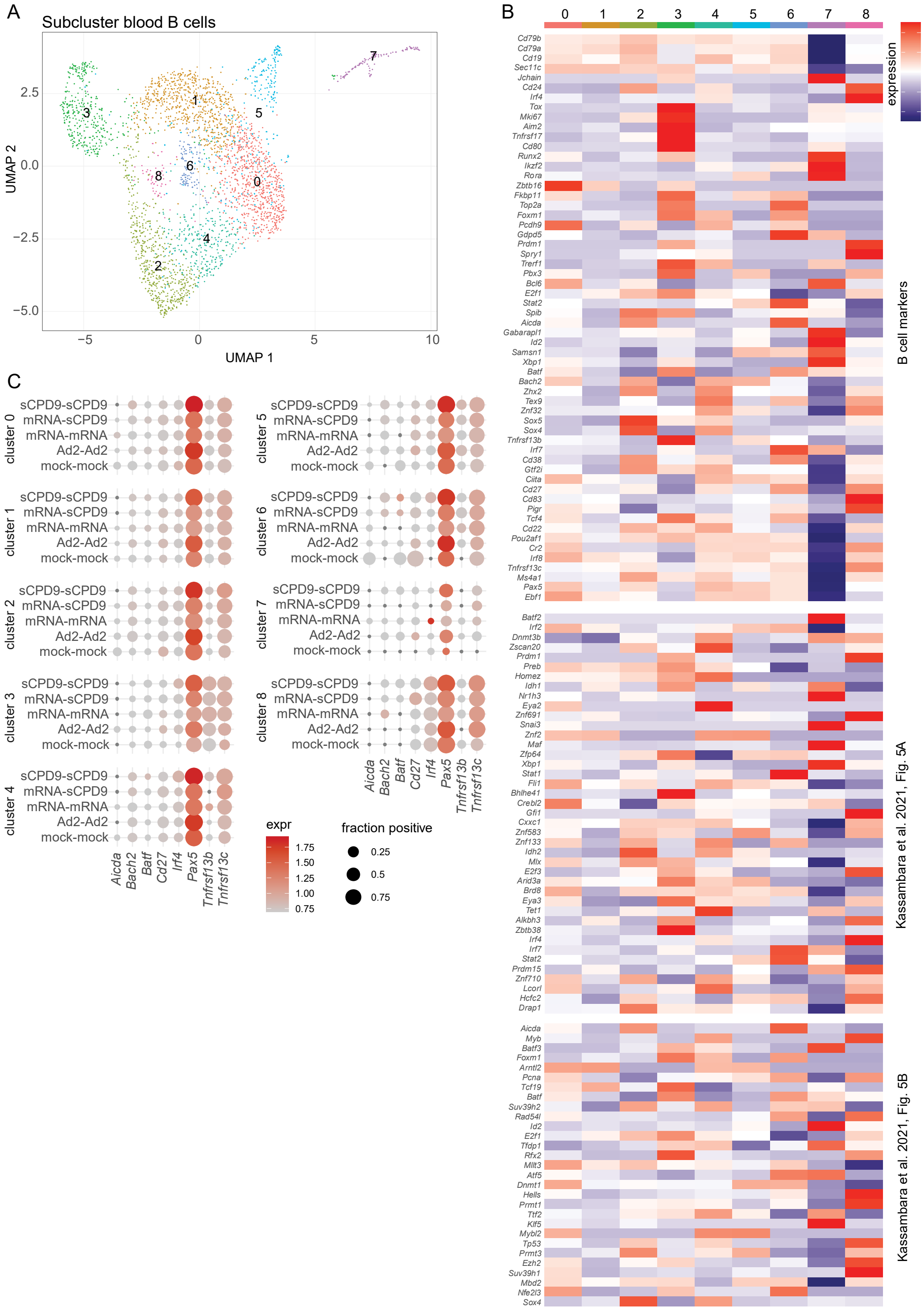


(**A**) Two-dimensional projections of single-cell transcriptomes using UMAP of blood B cells from prime-boost experiment. Cells are colored by clusters. Cluster 7 also contains cells expressing the T cell marker *Cd3e*, and therefore likely represents doublets or artefacts, and was therefore not considered further. (**B**) Heatmaps of classical B cell markers and genes from indicated publications. (**C**) Dotplots showing expression of selected marker genes in blood derived from B cell subcluster analysis. The size of the dot represents the fraction of cells in which at least one UMI of the respective was detected, the color is proportional to the average expression in those cells.

**Fig. S10.**

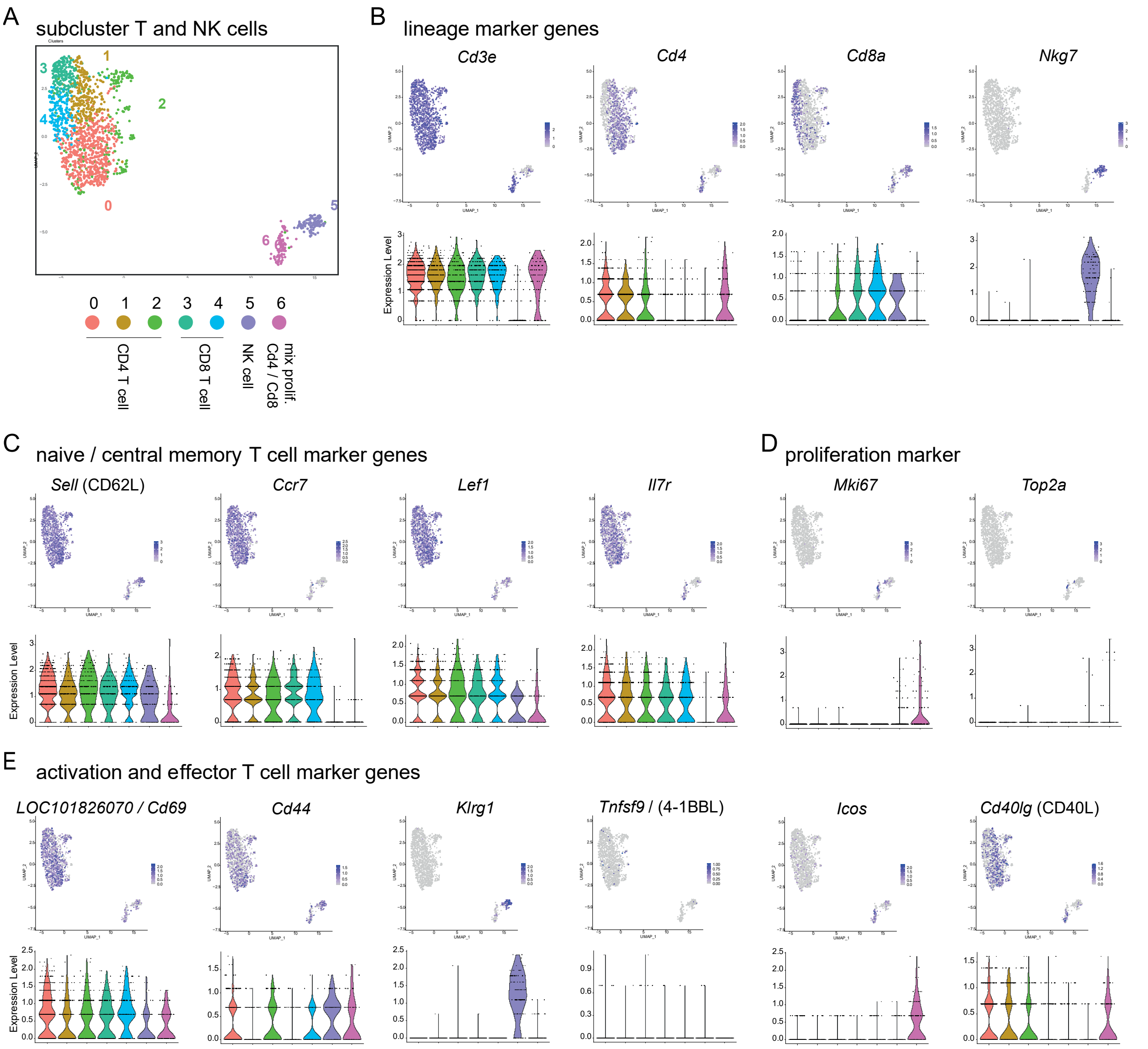


(**A**) Uniform manifold approximation and projection (UMAP) plot of blood T and NK cell sub-clustering from prime-boost experiment. Colors representing depicted cell types. (**B** – **E**) Feature plots showing gene expression patterns for individual cell markers and violin plots showing expression distributions in each cluster on single-cell level. Each dot represents a single cell. (**B**) Lineage marker genes: *Cd3e*, *Cd4*, *Cd8a*, *Nkg7*. (**C**) Naive / central memory T cell marker genes: *Sell* (CD62L), *Ccr7*, *Lef1*, *Il7r*. (**D**) Proliferation marker genes: *Mki67*, *Top2a*. (**E**) Activation and effector T cell marker genes: *LOC101826070* (*Cd69*), *Cd44*, *Klrg1*, *Tnfsf9* (4-1BBL), *Icos*, *Cd40lg* (CD40L).

**Fig. S11.**

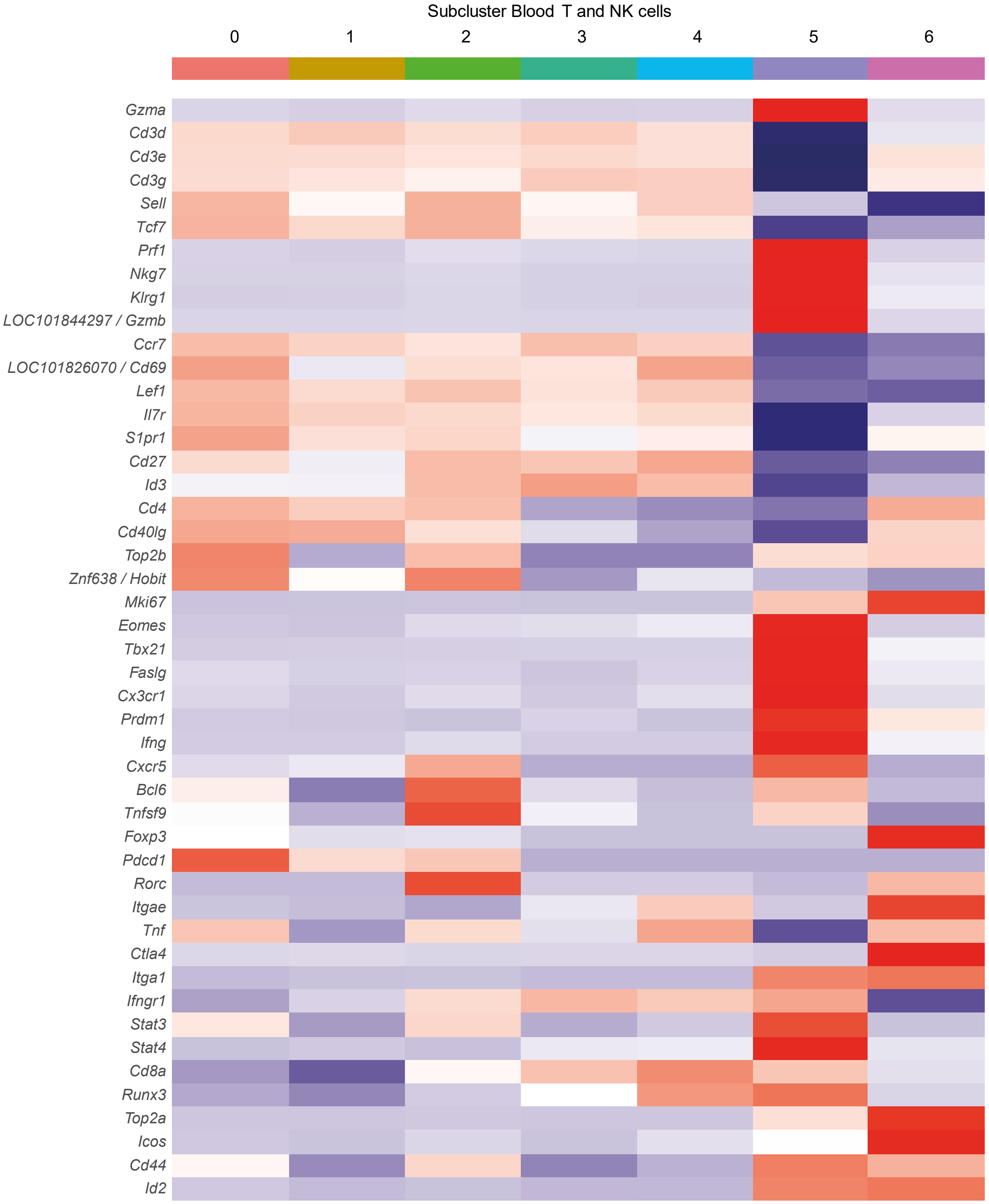


Heatmap of cell marker expression in identified clusters for blood T and NK cell subtypes from prime-boost experiment. Normalized average gene expression levels for cells in a cluster are indicated by coloration: low expression is shown in blue, high expression in red.

**Fig. S12.**

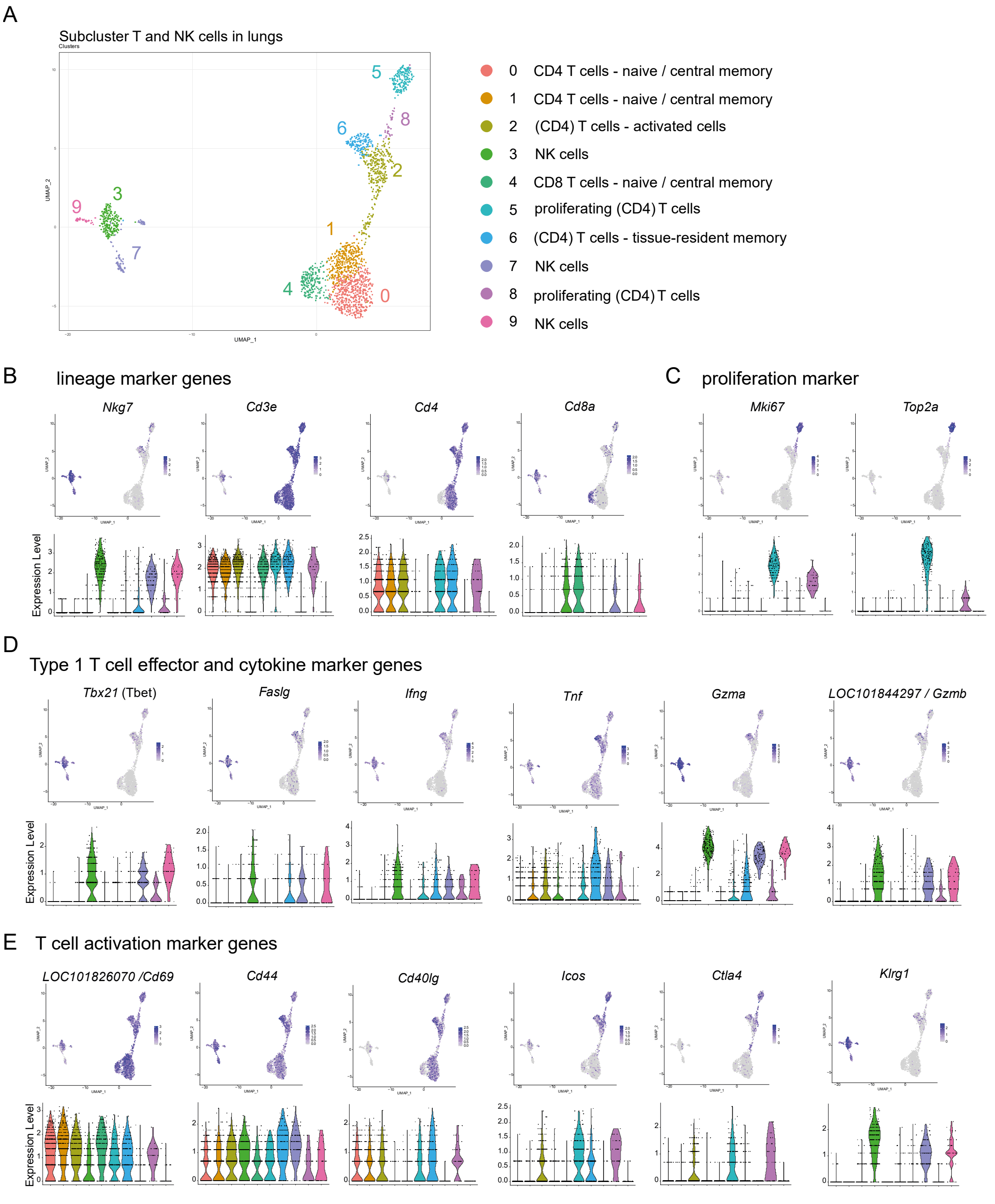


(**A**) Uniform manifold approximation and projection (UMAP) plot of lung T and NK cell sub-clustering from prime-boost experiment. Colours representing depicted cell types. (**B** – **E**) Feature plots showing gene expression patterns for individual cell markers and violin plots showing expression distributions in each cluster on single-cell level. Each dot represents a single cell. (**B**) Lineage marker genes: *Nkg7*, *Cd3e*, *Cd4*, *Cd8a*. (**C**) Proliferation marker genes: *Mki67*, *Top2a*. (**D**) Type 1 T cell effector and cytokine marker genes: *Tbx21* (Tbet), *Faslg*, *Ifng*, *Tnf*. (**E**) T cell activation marker genes: *LOC101826070* (*Cd69*), *Cd44*, *Cd40lg* (CD40L), *Icos*, *Ctla4*, *Klrg1*.

**Fig. S13.**

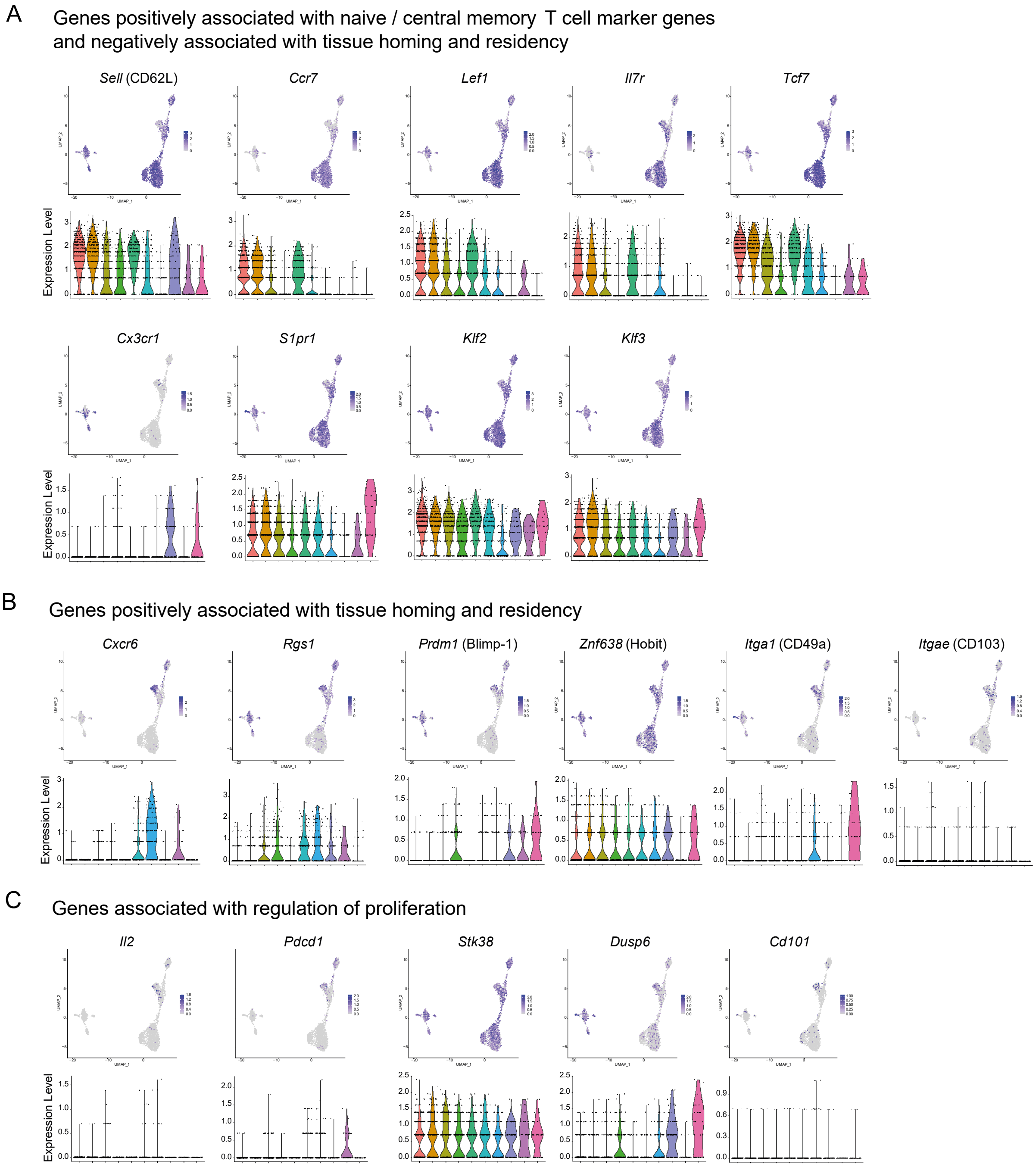


(**A** – **C**) Feature plots showing gene expression patterns for individual cell markers and violin plots showing expression distributions in each cluster on single-cell level from prime-boost experiment. Each dot represents a single cell. (**A**) Genes positively associated with naive / central memory T cell marker genes and negatively associated with tissue homing and residency: *Sell* (CD62L), *Ccr7*, *Lef1*, *Il7r*, *Tcf7*, *Cx3cr1*, *S1pr1*, *Klf2*, *Klf3*. (**B**) Genes positively associated with tissue homing and residency: *Cxcr6*, *Rgs1*, *Prdm1* (Blimp-1), *Znf638* (Hobit), *Itga1* (CD49a), *Itgae* (CD103). (**C**) Genes associated with regulation of proliferation: *Il2*, *Pdcd1*, *Stk38*, *Dusp6*, *Cd101*.

**Fig. S14.**

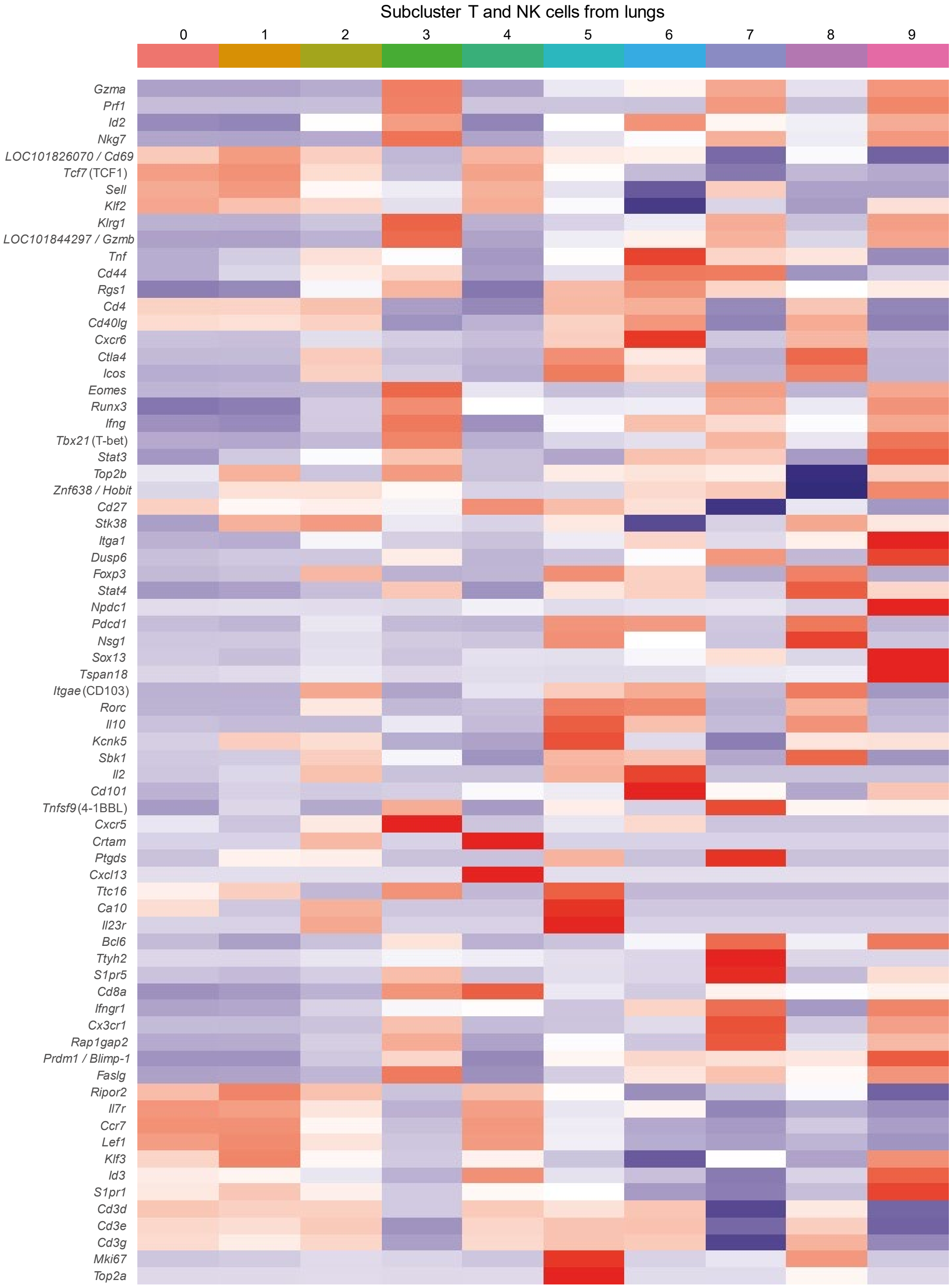


Heatmap of cell marker expression in identified clusters for lung T and NK cell subtypes from prime-boost experiment. Normalized average gene expression levels for cells in a cluster are indicated by coloration: low expression is shown in blue, high expression in red.

**Fig. S15.**

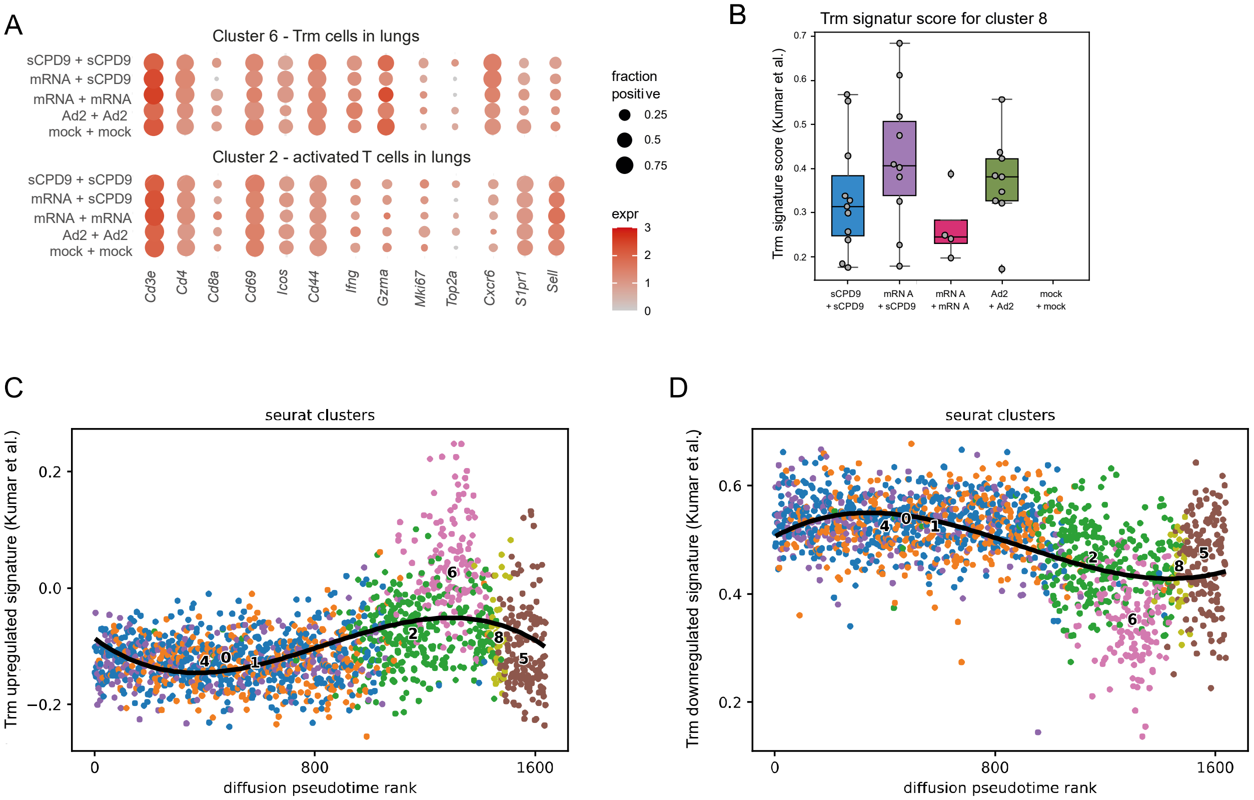


(**A** – **D**) Analysis of T cell subsets by scRNA-seq at 2 dpc in blood and lungs of prime-boost vaccinated hamsters. (**A**), Dotplots showing expression of selected T cell marker genes in the lungs in cluster 6 and 2 derived from T and NK subcluster analysis in Fig. S12A. The size of the dot represents the fraction of cells in which at least one UMI of the respective was detected, the color is proportional to the average expression in those cells. (**B**) Trm signature score (*41*) for individual cells in cluster 2 according to prime-boost vaccination regimen. Box, 25$^{th}$ to 75$^{th}$ percentile and whiskers Min to Max with individual cells displayed. (**C**) Trm upregulated signature (*41*) score as a function of diffusion pseudotime rank with black line showing a polynomial fit of degree three. (**D**) Trm downregulated signature (*41*) score as a function of diffusion pseudotime rank with black line showing a polynomial fit of degree three.

**Fig. S16.**

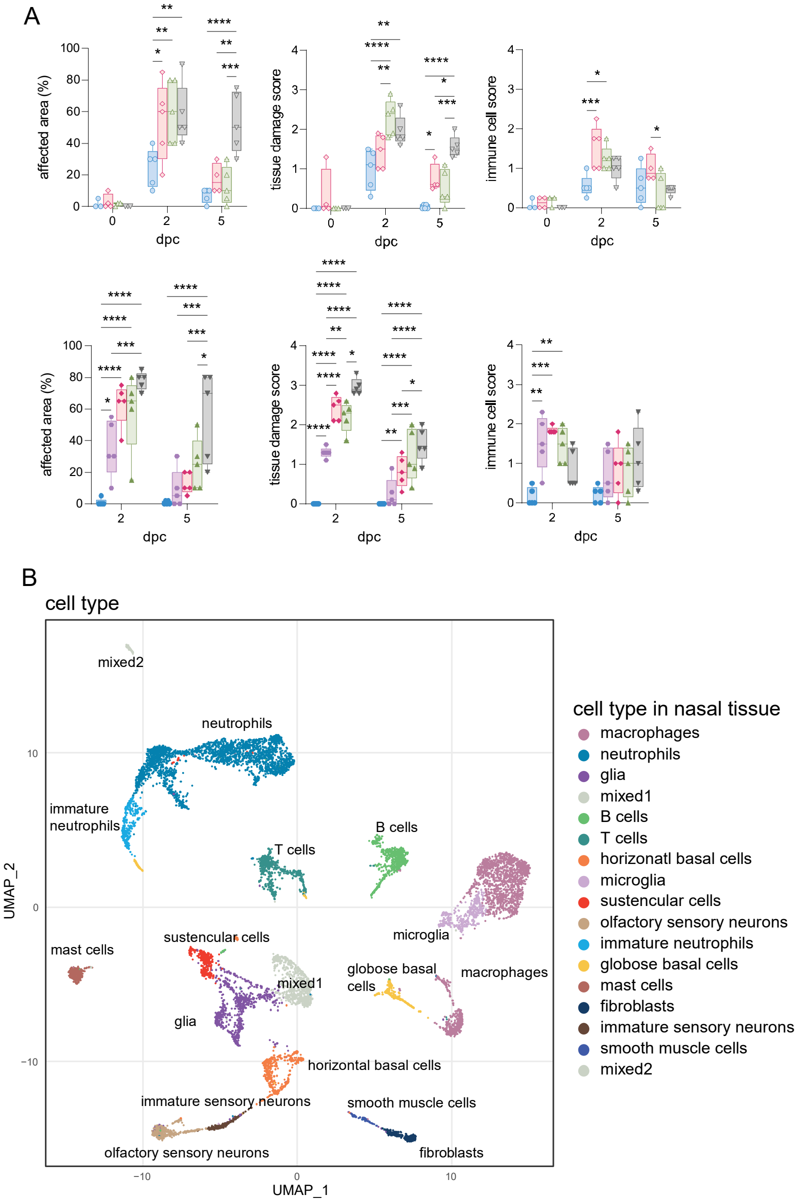


(**A**) Semi-quantitative scorings of histopathological findings in hamsters that received prime vaccination (top row) or prime-boost vaccination (bottom row). The area affected by mucosal damage due to SARS-CoV-2 infection is shown in percentage. The tissue damage score accounts for epithelial exfoliation, necrosis, apoptosis, cellular debris and cilium loss. Presence of neutrophils and lymphocyte in the nasal conchae is assessed by the immune cell score. Box (25th to 75th percentile) and whiskers (Min to Max). Two-way ANOVA and Tukey’s multiple comparison test were performed for statistical evaluation. $*p < 0.05$, $**p < 0.01$, $***p < 0.001$, $****p < 0.0001$. (**B**) Two-dimensional projections of single-cell transcriptomes using UMAP of nasal tissue cells from prime experiment. Coloration indicates cell types as annotated based on known marker genes.

**Fig. S17.**

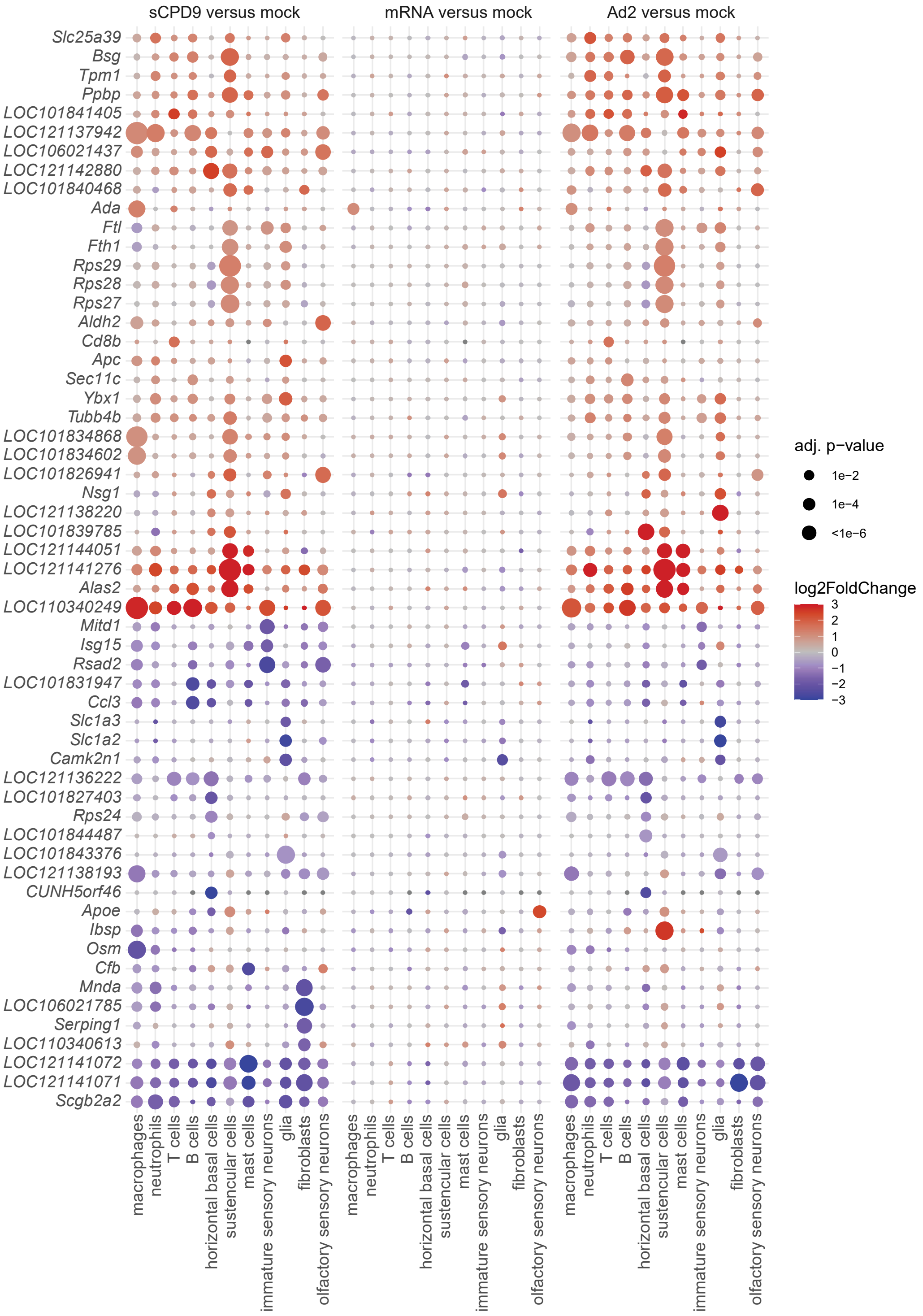


Dotplots showing fold changes of gene expression in indicated cell types of prime vaccinated compared to mock vaccinated animals. Coloration and point size indicate log2-transformed fold changes (FC) and p-values, respectively, in vaccinated compared to mock vaccinated animals. Adjusted (adj) p-values were calculated by DEseq2 using Benjamini–Hochberg corrections of two-sided Wald test p-values. Genes are ordered by unsupervised clustering.
